# Noncanonical P gene mRNA editing in Cedar virus generates a U protein that is required for efficient virion production

**DOI:** 10.64898/2026.02.27.708548

**Authors:** Henriette Schwotzer, Patricia Schupp, Richard Küchler, Dmitry S Ushakov, Gang Pei, Laura Rzepa, Sandra Diederich, Ilona Ronco, Olivier Reynard, Branka Horvat, Falk Butter, Stefan Finke

**Author notes:** Corresponding author: Stefan Finke. These authors contributed equally to the study (Henriette Schwotzer, Patricia Schupp).

## Abstract

Highly pathogenic Hendra and Nipah viruses encode accessory P gene products (C, V and W) that antagonize innate immunity and contribute to pathogenicity. Cedar virus (CedV), an apathogenic bat-borne henipavirus, is presumed to lack P gene mRNA editing and therefore is unable to express V and W proteins. Here, we identify CedV peptides originating from a frameshifted P gene open reading frame and demonstrate a previously unrecognized, noncanonical editing site at a homopolymeric adenine tract that introduces single-nucleotide A or G insertion. This mRNA editing produces a protein that we refer to as U protein, whose C-terminal domain shares sequence and predicted structural features with those of the henipavirus V protein. Recombinant CedV mutants defective in mRNA editing were only recoverable by trans-complementation and showed markedly reduced release of infectious virus in cell culture and attenuated replication in mice lacking type I interferon receptor. Our data revise the CedV gene expression models and reveal a noncanonical editing mechanism that supports the production of a U protein critical for efficient infectious virus release. These results expand the fundamental concepts of paramyxovirus gene expression and reveal an unexpected requirement for P-gene editing in efficient infectious-virus production, with implications for the evaluation of potentially high-consequence paramyxoviruses.

**Author Summary:** Cedar virus is a close relative of the highly pathogenic Nipah and Hendra viruses but is considered non-pathogenic. Unlike these viruses, Cedar virus was thought to lack a mechanism called P-gene mRNA editing, which allows related viruses to produce additional proteins that support infection. Here, we show that Cedar virus does use mRNA editing, but at an unusual sequence and by inserting either an adenine or a guanine nucleotide. This editing event generates a previously unknown protein, termed U, that shares selected features with the V and W proteins of other henipaviruses. Most strikingly, viruses unable to produce U released far fewer infectious virus, showed abnormal membrane-associated structures, and replicated less efficiently in susceptible mice. These findings revise the current model of Cedar virus gene expression and reveal that mRNA editing can contribute directly to efficient virus production, not only to immune evasion. More broadly, our results highlight the need to search for unconventional editing sites when annotating and assessing newly discovered paramyxoviruses.

## Introduction

Henipaviruses (genus *Henipavirus*, family *Paramyxoviridae*) include the biosafety level 4 (BSL-4) pathogens Nipah virus (NiV) and Hendra virus (HeV). They are enveloped, single-stranded, negative-sense RNA viruses whose genomes encode nine proteins. Three accessory P-gene products (C, V and W) are major pathogenicity factors. The C protein is translated from an alternative downstream open reading frame by leaky scanning, whereas the V and W proteins arise from cotranscriptional mRNA editing that inserts one (in the case of V) or two (in the case of W) guanine nucleotides in the nascent mRNA. This causes a frameshift and generates these distinct C-terminal variants of P (reviewed in [1]).

Cotranscriptional P gene editing is broadly conserved among paramyxoviruses and was first inferred from C-terminally distinct V proteins in simian virus 5 [2]. V proteins typically contain a conserved cysteine-rich zinc-finger domain, although truncations without this protein domain have been proposed in some paramyxoviruses, such as Jeilongvirus. Exceptions to V protein expression have indeed been reported in human parainfluenza virus type 1 and Cedar virus (CedV) [3], where the canonical editing motif is absent [4, 5].

In NiV and HeV, the proteins C, V and W antagonize innate immune signaling by interfering with pattern recognition receptor pathways and inhibiting STAT1/2□mediated interferon responses [6–10]. Additionally, protein W can impair NF□κB signaling through 14□3□3 binding and nuclear sequestration of p65 [11], and it has been linked to the modulation of metabolism and apoptosis [12]. These immune evasion strategies are thought to promote sustained replication in immune-competent respiratory epithelia [13].

CedV is a Pteropus bat-associated henipavirus that is considered apathogenic. CedV was reported to lack the conserved P gene editing motif (UUUUCCC, genome template orientation) present in NiV and HeV and to harbor a premature stop codon in the putative V open reading frame, suggesting an inability to express functional V. The presumed absence of V and W expression has been proposed to underlie the non-pathogenic phenotype of CedV [4]. Furthermore, the lower STAT1 binding capacity of CedV P than that of NiV and HeV [14] and the difference in Ephrin receptor usage between CedV and pathogenic henipaviruses [15, 16] are considered important for the different pathogenic potential of CedV.

Despite this, CedV can replicate and disseminate to limited levels in vivo [4, 17, 18] in the presence of an intact type I interferon response. Furthermore, CedV circulates in Pteropus bat colonies, indicating sufficient replication for maintenance in reservoir populations.

Here, we use proteomics, targeted sequencing and reverse genetics to identify a previously unrecognized CedV P gene product generated by non-canonical mRNA editing. We show that editing yields a U protein that shares features of the henipavirus V and W proteins and that disruption of this editing event severely impairs the production of infectious viruses both in vitro and in vivo. These findings revise CedV gene annotation and reveal atypical editing mechanisms in henipaviruses that support CedV replication.

## Results

### Identification of U peptides in the CedV infection proteome

Based on our recent observation that SK-N-SH neuroblastoma cells are highly permissive to CedV infection and support robust virus production, we selected this cell line for an unbiased Proteomic analysis of cell lysates from recombinant CedV (rCedV)-infected SK-N-SH cells at 48 hours post-infection (hpi) revealed the expression of all the canonical viral proteins (N, P, M, F, G, L and C) (Fig. 1A). To detect previously unknown proteins, we additionally included an in silico translation of all possible ORFs with at least 20 amino acids from the CedV genome during peptide identification. Notably, we found three additional peptides that mapped to the same alternative reading frame within the P gene (genomic nt 3308–3517), corresponding to a stretch of 70 amino acids translated from frame 3 of the P gene (Fig. 1C-E). Two of these peptides showed 44–48% sequence identity to a consensus sequence derived from the C-terminal domains of V proteins from NiV and HeV, indicating the possibility of a U protein with some but limited similarities to henipavirus V proteins in CedV (Fig. 1G). Proteome analysis in PTAGV cells derived from *Pteropus giganteus* bats (B) confirmed the presence of peptide p2 and thus translation of the alternative P gene product (Fig. 1F). Alignment of the putative unique C-terminal domain (CTD) of the CedV U protein with the NiV V protein CTD revealed partial sequence homology within peptide 3, whereas no homology was observed in the flanking upstream or downstream regions (Fig. 1H).

**Figure 1:**
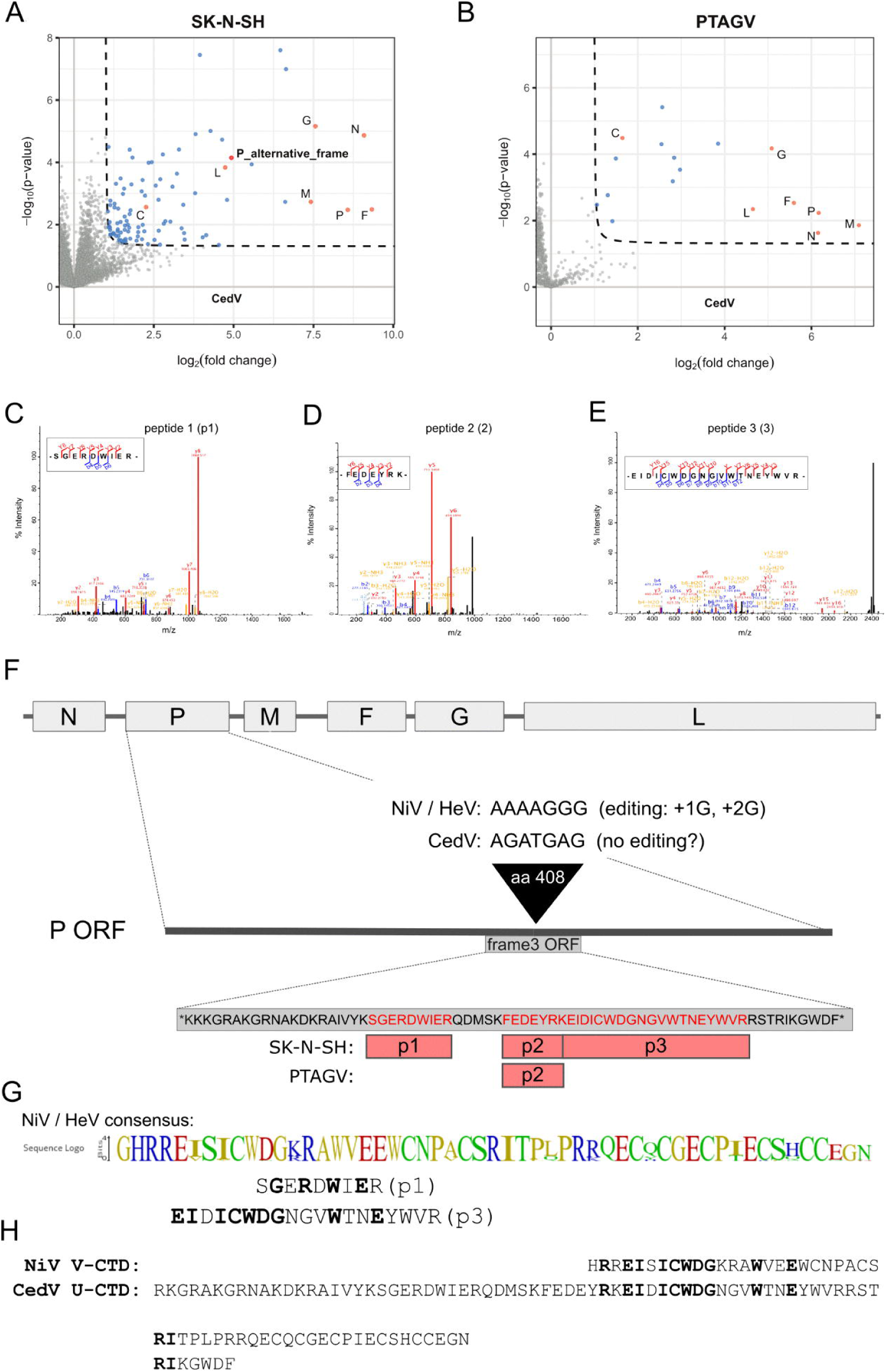
Identification of alternative P gene reading frame peptides in CedV-infected cells and their similarity to those of the henipavirus V protein. (A) Volcano plot showing upregulated proteins in CedV-infected SK-N-SH cells, as determined by label-free quantitative proteomics (n = 4). Enriched proteins (p value < 0.05 (Welch’s t test) and fold change (FC) > 2) are marked in blue. Canonical CedV proteins are highlighted in orange, and the CedV P gene alternative frame is highlighted in red. (B) Comparable Volcano plot for CedV infected bat PTAGV cells (C–E) Representative MS/MS spectra of the three detected peptides with b and y ions. (F). Schematic drawing of the CedV genome with annotated ORFs and the nucleotide sequence around amino acid position 408 of the P gene with the editing site of NiV and HeV. The localization of the identified peptides to the 70-amino acid stretch of ORF frame 3 is illustrated for the SK-N-SH and PTAGV cell proteome analysis. (G) Consensus sequence generated of the C-terminal domains of the HeV and NiV V proteins on the basis of 57 gene bank sequences (list of sequences at DOI 10.5281/zenodo.18642052). Peptides p1 and p3 mapped to the consensus sequence, whereas p2 did not show similarity to the consensus sequence. Positions with amino acid identity between the consensus sequence and CedV peptide are marked by bold letters. (H) Alignment of the NiV V and CedV U CTDs. Identical amino acids in V and U are marked by bold letters.

### A noncanonical mRNA editing site generates CedV U transcripts

To test whether these peptides arise from mRNA editing, we subjected RT-PCR amplicons from infected human SK-N-SH and HeLa cell lines and bat PATGV cells to next-generation sequencing (NGS). Bioinformatic analysis revealed a hotspot of single-nucleotide insertions at genome position 3312 (P-mRNA position 1299). Unlike canonical paramyxovirus editing, insertions comprising either adenine or guanine nucleotides were detected using both oligo(dT) and negative-sense RT primers across different cell types at frequencies of 13 to 20% (Fig. 2A, S8 Table 2). The insertion site comprises a heptameric adenine stretch (TTAAAAAAAGAAA, mRNA orientation) and is located upstream of the canonical editing sites in NiV and HeV. Cloning and Sanger sequencing of independent cDNA clones confirmed the insertions and supported genuine editing rather than sequencing artifacts.

**Figure 2:**
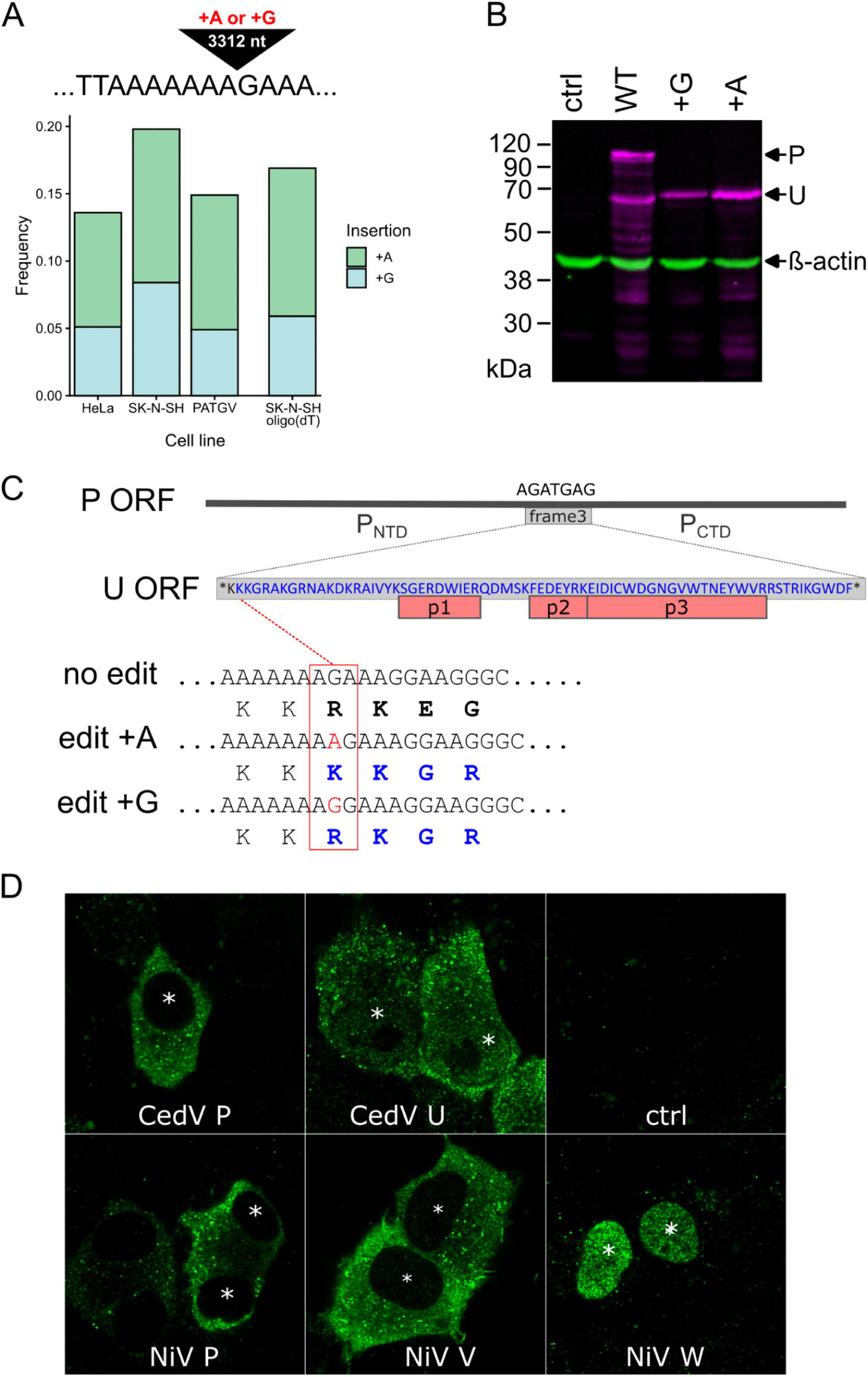
Positions of CedV mRNA editing, expression and localization of U proteins. (**A**) Proportion of editing events detected in P gene mRNA amplified by either negative (-) sense or oligo-dT reverse transcription (RT) primers from isolated total RNA from CedV-infected SK-N-SH, HeLa, and PATGV cells. Frequencies of A and G insertions [1 =100%] are indicated by green and blue, respectively. **(B)** Western blot of Strep-tagged P variants in BSR-T7/5 cells expressed from the cloned and mutated P gene cDNA sequences at 20 h post transfection. **(C)** Insertions at position 3312 (red letters) induce a frameshift at the second codon of the frame 3 70-aa ORF, leading to the expression of a 69-aa alternative C-terminal domain (CTD; blue letters) containing the identified peptides 1, 2, and 3.U **(D)** Intracellular localization of expressed Strep-tagged CedV and NiV P protein variants expressed from their cDNA sequences in BSR-T7/5 cells at 20 h post transfection. Ctrl: BSR-T7/5 cells transfected with the empty vector. The nuclei are marked by a white star.

### Editing-dependent expression and localization of a CedV U protein

cDNAs from the P gene with single adenosine or guanosine nucleotide insertions at the identified editing site were transfected into BSR T7/5 cells, leading to the synthesis of lower-molecular-weight U proteins (Fig. 2B). The A or G nucleotide insertions at the noncanonical site introduce a frameshift and permit translation of a 69-amino-acid alternative C-terminus that encompasses the proteomics-identified peptides (Fig. 2C). Compared with NiV P and V, the CedV P protein remained predominantly cytoplasmic in transfected cells, whereas the V-like protein localized to both the cytoplasm and the nucleus (Fig. 2D), suggesting that the alternative C-terminus provides additional trafficking determinants and may combine the properties of the NiV V and W proteins.

### Structural and motif predictions identify selected features shared with henipavirus V and W proteins

AlphaFold predictions indicated that the CedV U protein had a largely disordered architecture with short structured elements and a characteristic two-β-sheet protrusion in the C-terminal domain (CTD), resembling the NiV V protein CTD (Fig. 3A, S2 Figure 1). As expected, the global pTM (predicted template modeling) scores were low for both CedV U and NiV V (0.22 and 0.17, respectively), reflecting the extensive disorder predicted across the full-length proteins. In contrast, the residue-level pLDDT scores indicated high confidence in the predicted structures of the C-terminal domains (S2 Figure 1). Whereas the ß-sheet structure was highly similar between CedV and NiV, CedV mRNA editing led to the formation of a positively charged stretch of five amino acids that is not present in the NiV V protein (Fig. 3B). Notably, while the NiV V CTD structure was identically predicted also when only the CTD was used, for the 2 beta-sheet fold of CedV U CTD the complete U protein was required, indicating a role of the N-terminal region in CTD folding.

**Figure 3:**
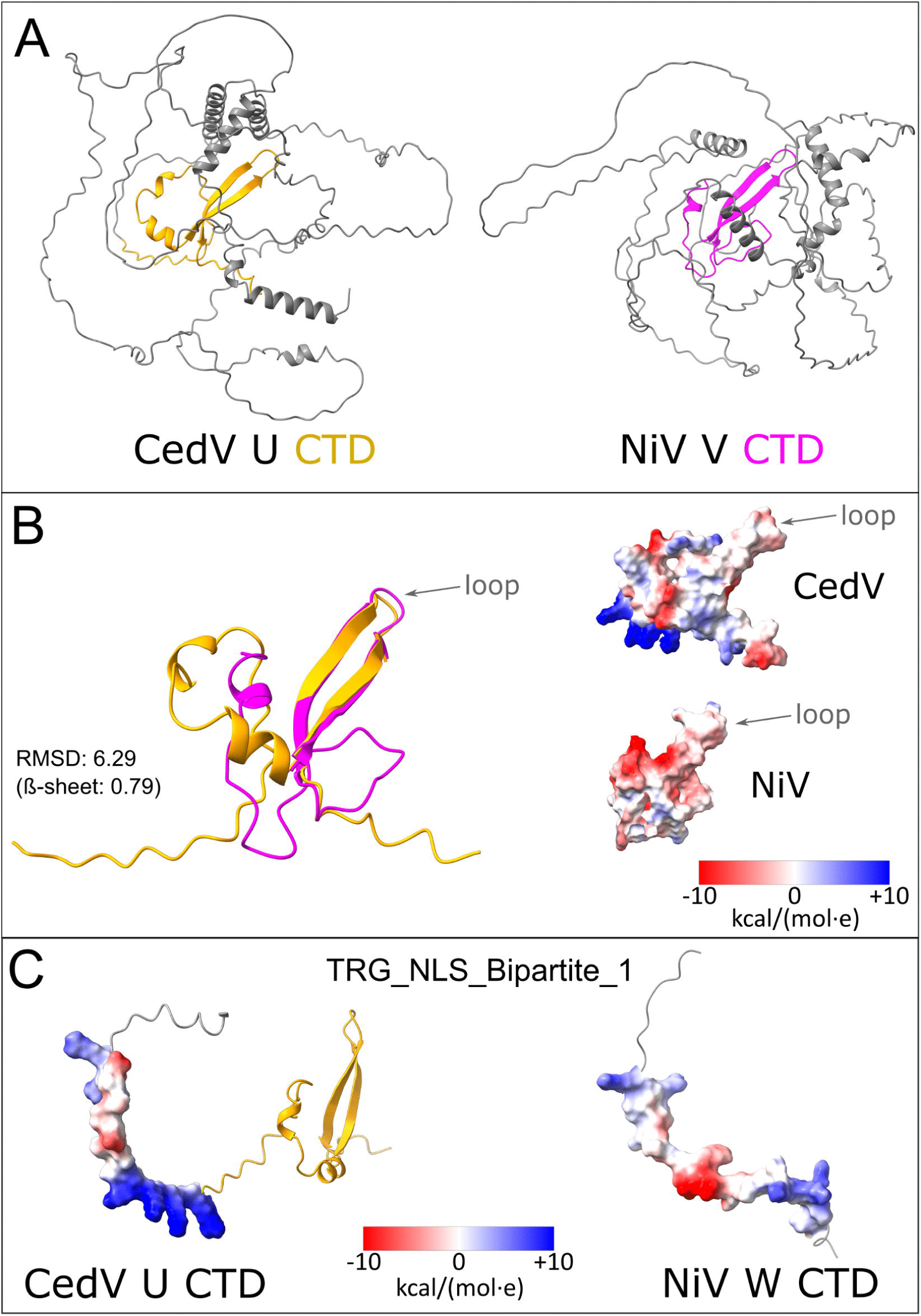
Structural predictions of the CedV U and NiV V proteins. **(A)** Alphafold prediction of the CedV U and NiV-V structures with CTDs labeled in orange and magenta, respectively. Residue-resolved pLDDT scores are provided in S1 Figure. **(B)** Ribbon diagrams of CedV U and NiV V CTD overlays. RMSD: Root mean square deviation. Right: space filling representation of the CTDs with an applied electrostatic potential surface. Blue: positive charge, red: negative charge. The ß-sheets connecting loops are indicated. **(C)** Predicted bipartite nuclear localization signal (NLS) in the CedV U protein, which consists of positively charged amino acids upstream of the editing site and its CTD. A comparable bipartite NLS in the NiV W protein is generated by editing.

Linear-motif searches using ELM [19] predicted candidate interaction motifs (including a putative 14-3-3 binding motif) (S3 Figure 2) and a bipartite nuclear localization signal created by mRNA editing (Fig. 3CD). These features are consistent with nuclear localization and potential interactions with host factors. Notably, a potential binding site for the kinesin light chain (Lig_KLC1S3WD_1) was found in the ß-sheets connecting the loops of the CedV U and NiV V proteins (S3 Figure 2). However, neither a bipartite NLS nor a 14-3-3 binding motif was found in the NiV V protein. Instead, these features are part of the NiV W protein, which contains a bipartite NLS at the editing site and two 14-3-3 binding motifs in its CTD (Fig. 3C).

The functional relevance of the predicted nuclear localization signal (NLS) was assessed using two U mutants, U-NLSmut-I and U-NLSmut-II. Expression of either mutant significantly reduced nuclear localization to levels comparable to those observed for P, although a residual fraction of U remained detectable in the nucleus (Fig. 4BC). Further evidence for nuclear shuttling of U was obtained by inhibiting exportin 1-mediated nuclear export with leptomycin B (Figure 4D; S4 Figure 3). This treatment caused pronounced nuclear accumulation of U. U-NLSmut-I also accumulated in the nucleus, but to a significantly lesser extent than wild-type U, supporting an important role of the predicted NLS in the nuclear shuttling of U. In contrast, P showed no detectable nuclear accumulation even after leptomycin B treatment, indicating that nuclear shuttling is a specific property of the editing product U.

**Figure 4.**
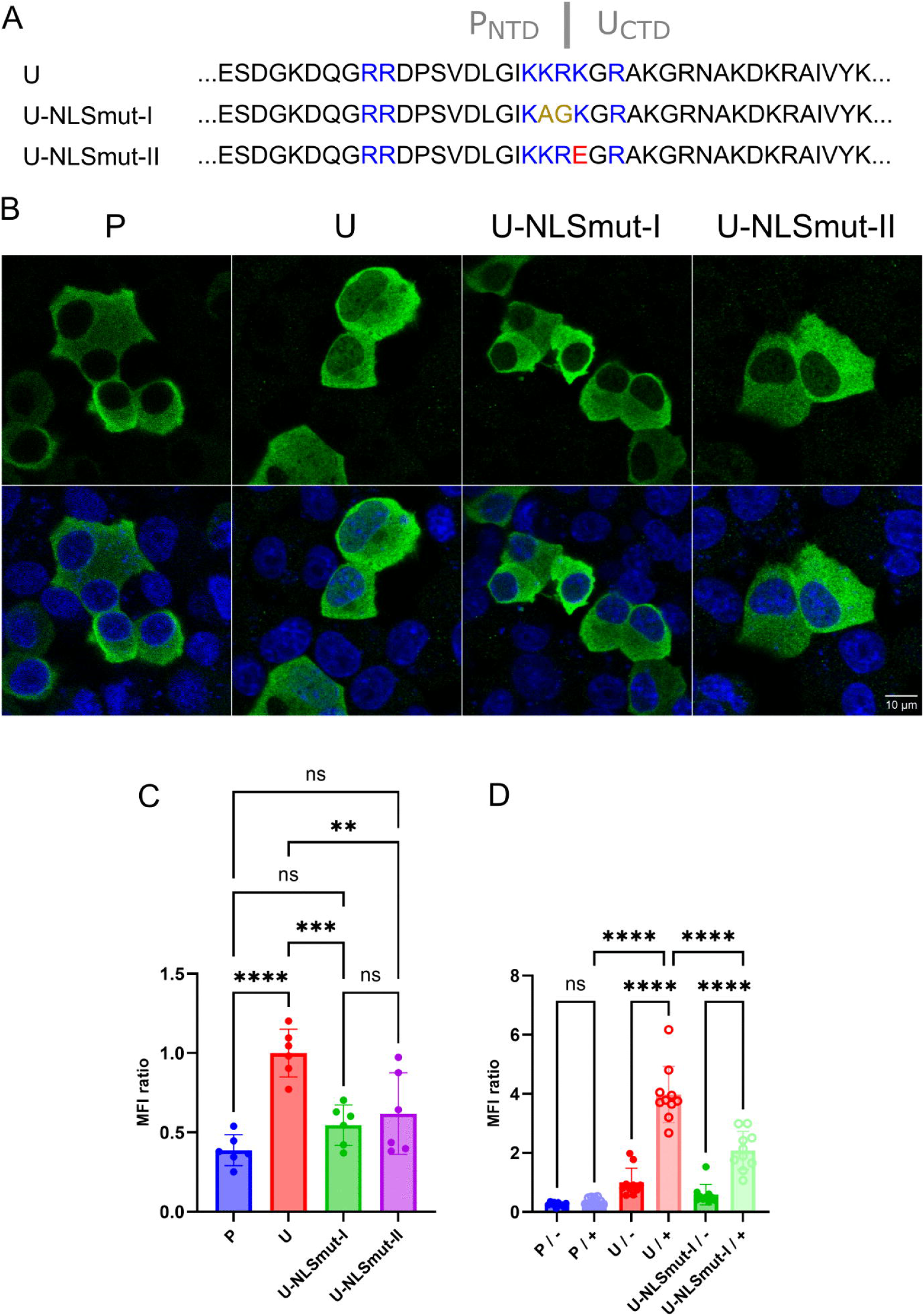
Nuclear localization of U depends on a predicted nuclear localization signal and nucleocytoplasmic shuttling. (**A**) Positively charged residues within the predicted bipartite nuclear localization signal (NLS) in the PNTD and UCTD are highlighted in blue. U-NLSmut-I contains amino acid substitutions at positions 400 and 401, in which the positively charged residues KR were replaced with the neutral residues AG. In U-NLSmut-II, the positively charged lysine residue at position 402 was replaced with the negatively charged glutamate residue (K402E). **(B)** Immunofluorescence detection of Strep-tagged P, U, and the indicated mutants in BSR-T7/5 cells at 20 h post-transfection. **(C)** Quantification of the nuclear localization of the indicated proteins. Nuclear mean fluorescence intensities (MFIs) were normalized (the mean nuclear MFI was set to 1) and are presented as nuclear-to-cytoplasmic MFI ratios. Statistical significance is indicated as follows: *P < 0.05, **P < 0.01, ***P < 0.001,****P < 0.0001, ns, not significant. **(D)** Impact of leptomycin treatment on nuclear localization of U, U-NLSmut-I and P.

Structural and linear motif predictions together with functional data related to an editing-dependent nuclear localization signal indicated that the CTD of the CedV U protein shares the typical features of both the NiV V protein and the W protein.

### Distinct localization of P-and U proteins in CedV-infected cells

Upon transfection, the CedV P protein accumulated in viral cytoplasmic inclusion bodies (IB) in CedV-infected cells, whereas the U protein was only marginally enriched in IBs (Fig. 5AB). Instead, the U protein remained largely diffuse in the cytoplasm and nucleus, indicating different subcellular distributions of P and U protein during infection and supporting a role for the CTD of P in recruitment to IBs. Notably, U protein displayed prominent nuclear localization in both infected (Fig. 5AB) and noninfected cells (Fig. 5CD).

**Figure 5:**
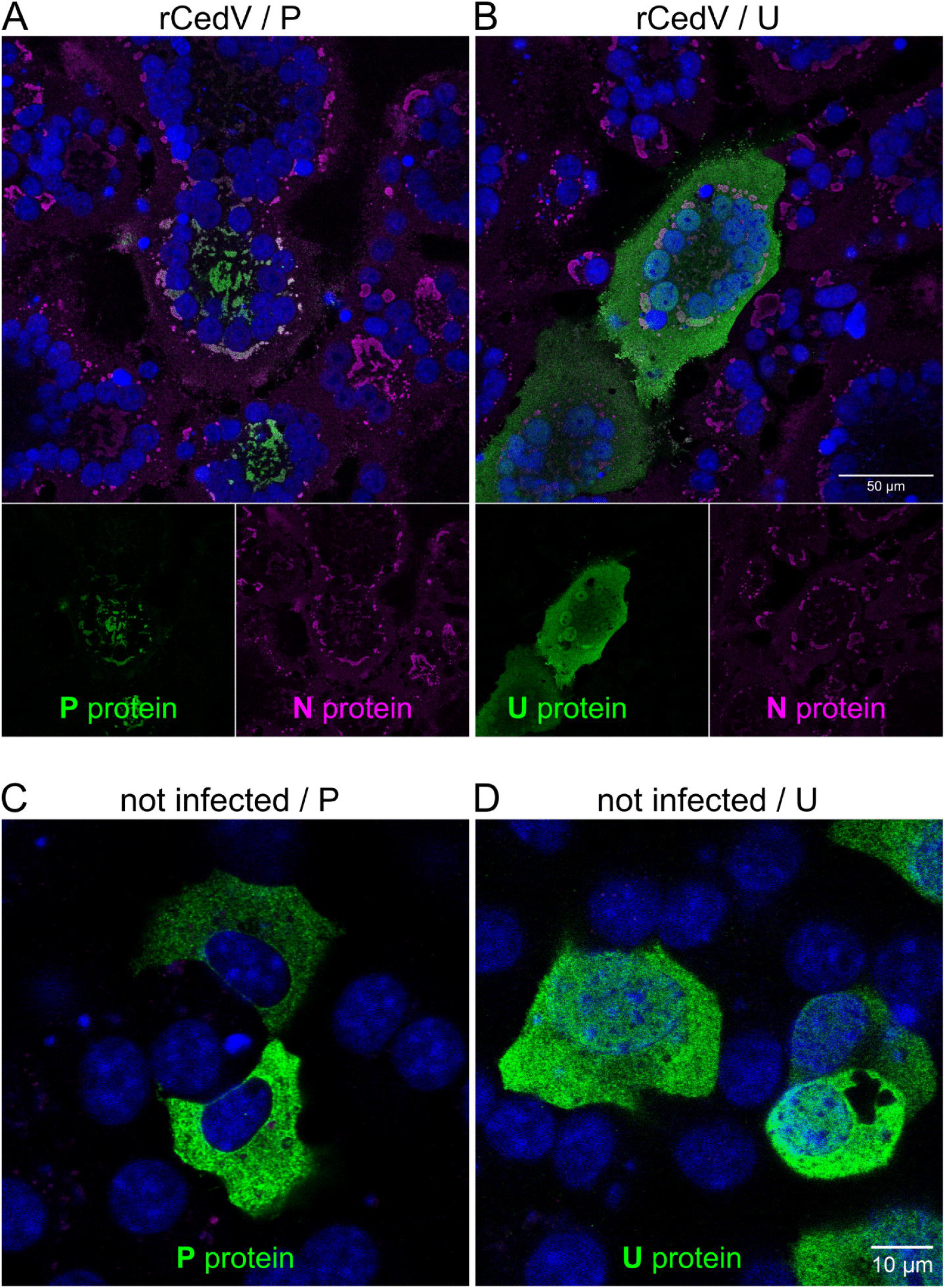
Immunofluorescence detection of Strep-tagged CedV P and U proteins in rCedV-infected and noninfected cells. (**A, B**) Two hours after infection (MOI = 0.1), BSR-T7/5 cells were transfected with plasmids encoding N-terminally Strep-tagged CedV P or U proteins. Immunofluorescence staining was performed 16 h posttransfection with antibodies specific for the Streptag (green) and CedV N proteins (magenta). **(C, D)** Plasmid-derived CedV P and U protein localization in noninfected cells. Blue: Hoechst 33324 stain.

### CedV U protein suppresses type I interferon induction but not interferon signaling

Given that antagonism of innate immune pathways is a hallmark of henipavirus V and W proteins, we compared the ability of the CedV U protein to inhibit type I interferon induction and downstream interferon responses with that of known regulators such as the CedV P and the NiV P, V and W proteins. Upon stimulation of RIG-I-or TBK1-dependent signaling, expression of the U protein reduced IFN-ß promoter–driven reporter activity. While the antagonistic activity of the CedU protein was significantly lower than that of the NiV V protein after RIG-I-mediated stimulation, no significant differences were observed after TBK-1-mediated induction (Fig. 6AB). In contrast, whereas all three tested NiV proteins potently inhibited IFN-induced signaling from approximately 7-fold to 13-fold, CedV P only modestly reduced luciferase reporter activity by 3-fold, and the CedV U protein did not significantly suppress the interferon response (Fig. 6C). Western blot analyses confirmed molecular weights and comparable expression of the tested virus proteins (Fig. 6D).

**Figure 6:**
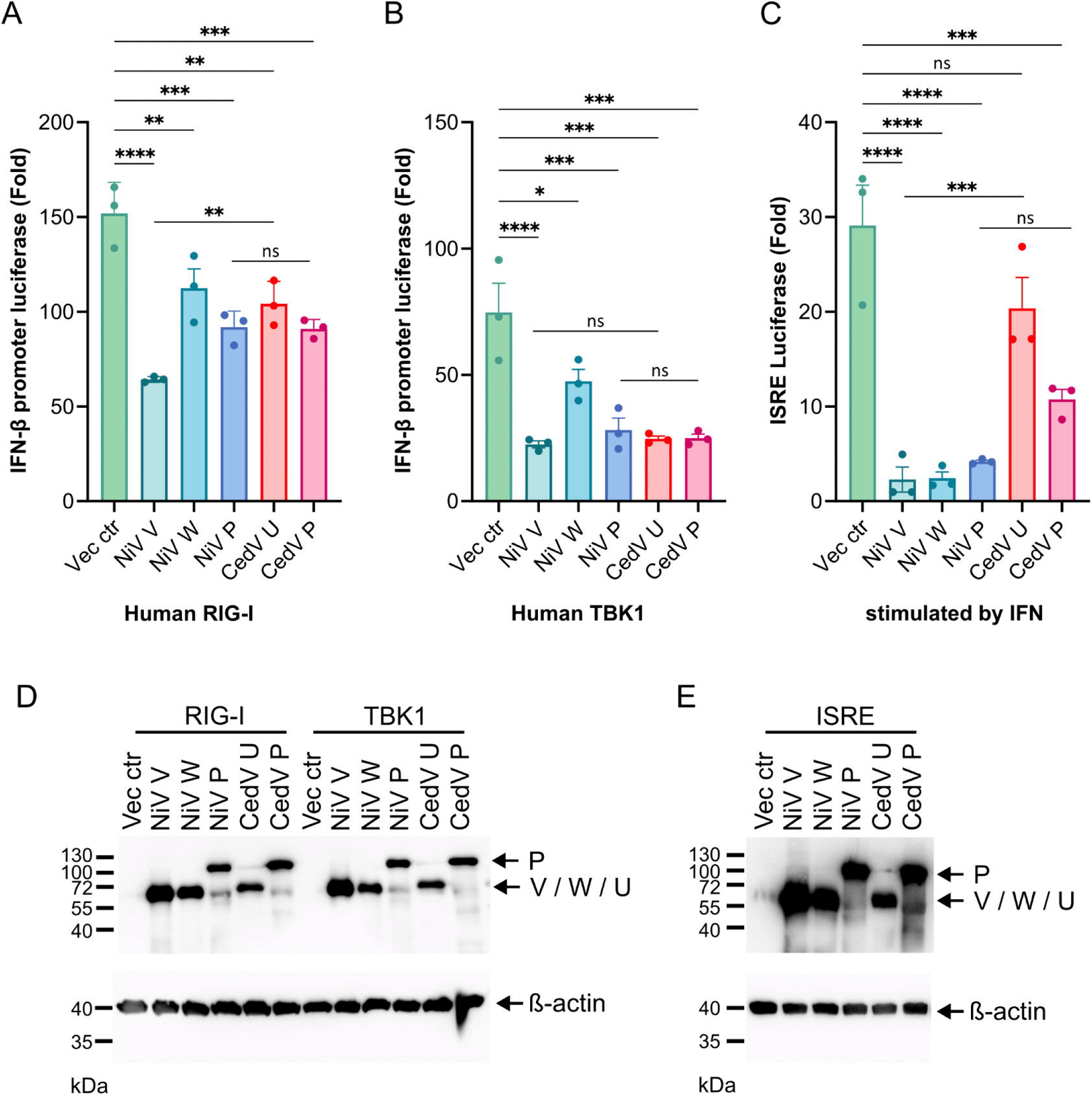
CedV U protein inhibits IFN induction but fails to inhibit the interferon response. Inhibition of IFN-ß promoter induction in HEK293T cells expressing the indicated NiV and CedV P gene products together with human RIG-I **(A)** and TBK1 **(B)** at 24 h post transfection. **(C)** Interferon-stimulated response element (ISRE) activation was measured in HEK293T cells expressing the indicated NiV and CedV P gene products upon universal IFN-I stimulation for 16–24 h. Data are presented as the mean ± SEM. Statistical analysis was performed using one-way ANOVA, followed by the Dunnett multiple comparison test. P values are indicated as ns (*p* > 0.05), * (*p* ≤ 0.05), and ** (*p* ≤ 0.01). *** (*p* ≤ 0.001) and **** (*p* ≤ 0.0001). Each dot represents one independent experiment. (D,E) Western blot detection of P gene products and cellular ß-actin.

### Generation of recombinant, editing-defective CedV

To validate the newly identified editing site, we disrupted the adenine heptamer immediately upstream of the guanine at position 3312 by substituting two of the adenine residues with guanines. In addition, we introduced synonymous marker mutations in the P open reading frame, enabling the detection of potential rapid accumulation of mutations and restoration of a functional editing site following virus rescue (Fig. 7A).

**Figure 7:**
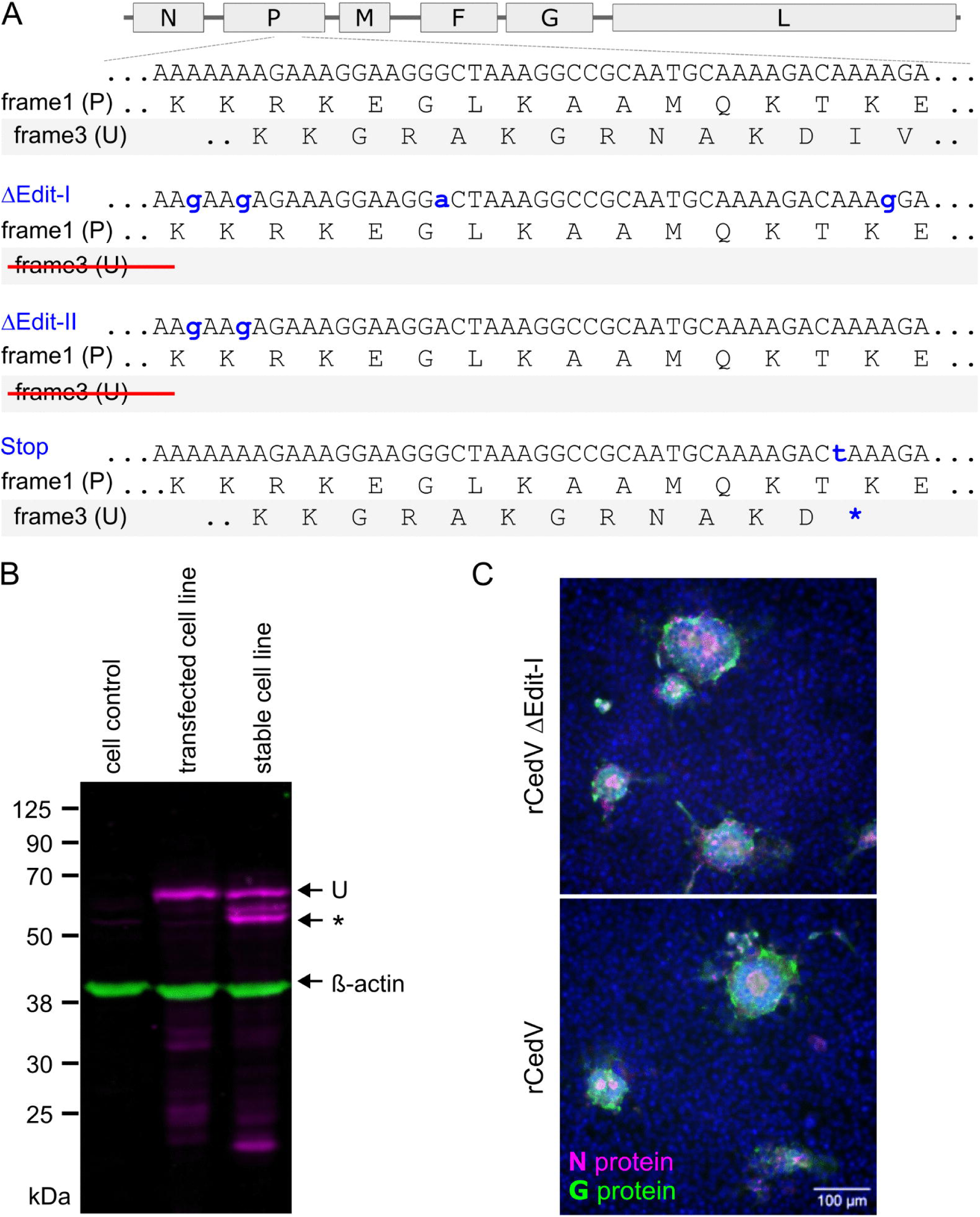
Rescue of mRNA editing-defective rCedV ΔEdit in cells complementing U protein. **(A)** Disruption of the heptameric adenine stretch of the CedV editing site, as indicated by blue lowercase letters. With respect to rCedV-ΔEdit-I, downstream guanine and adenine stretches were altered by synonymous mutations in the P protein. For rCedV ΔEdit-II, only the homopolymeric adenine tract of the editing site was modified. For rCedV-Stop, generation of a stop codon in the U ORF at the marked position in blue resulted in a truncated U protein with only 12 amino acids C-terminally of the editing site. **(B)** Western blot detection of the Strep-tagged CedV U protein in expression plasmid-transfected BSR-T7/5 cells and in the cell line stably expressing the U protein BSR-T7-StrepU. *: presumably truncated U protein. **(C)** Infection of noncomplementing BSR-T7/5 cells with rCedV ΔEdit-I after virus rescue in U complementing BSR-T7-StrepU cells and rCedV at 22 hpi (magenta: N protein, green: G protein, blue: Hoechst 33324 stain).

Despite multiple rescue attempts, rCedV ΔEdit-I could not be recovered from the cDNA plasmid, although all changes were synonymous for protein P. This rescue failure demonstrates an essential function provided by the U protein product that can be expressed only with a functional editing site. We presumed that this failure could be overcome by trans-complementation with ectopically expressed U protein.

To complement the editing-defective genome, we generated BSR-T7-StrepU cells stably expressing an N-terminally Strep-tagged CedV U protein. Expression was confirmed by Western blotting (Fig. 7B). Rescue by trans-complementation on BSR-T7-StrepU cells was successful, as demonstrated by the transfer of infectious virus via culture supernatants to noncomplementing BSR-T7/5 cells.

Notably, infection of noncomplementing cells with complemented virus resulted in N and G protein expression and syncytia formation comparable to those of wild-type rCedV, indicating that U protein expression is not required for viral gene expression or cell–cell fusion (Fig. 7C). Deep sequencing of the supernatants from two independent rescue experiments confirmed the presence of the engineered rCedV ΔEdit-I sequence.

Similarly, rCedV ΔEdit-II (disruption of the A-heptamer only) and rCedV-Stop (a stop codon in the ORF of the U C-terminal domain; Fig. 7A) were also only recovered in trans-complementing BSR-T7-dStrepU cells, and their sequences were also unchanged, as verified by NGS.

Amplicon sequencing of the P gene mRNA from rCedV ΔEdit-I and rCedV ΔEdit-II revealed a complete loss of editing in both mutants (Fig. 7A, S9 Table 3). In contrast, an editing frequency of 27% in rCedV-Stop confirmed that P gene mRNA editing was maintained in this virus mutant. These results functionally validate the requirement of the homopolymeric seven adenine tract for CedV P gene mRNA editing. Importantly, the availability of editing-negative mutants without any amino acid changes in the P protein enabled us to specifically assess how P gene editing and expression of the corresponding U protein contribute to CedV replication.

### The expression of U proteins is required for the efficient release of infectious virus particles

Whereas wild-type rCedV infection spread markedly from day 1 to day 2 and produced multiple small secondary syncytia, two independently rescued editing-defective clones (rCedV ΔEdit-Ia and rCedV ΔEdit-Ib) formed almost exclusively large syncytia (Fig. 8B). This pattern is consistent with reduced dissemination via cell-free viruses and a shift toward spread by cell–cell fusion.

**Figure 8:**
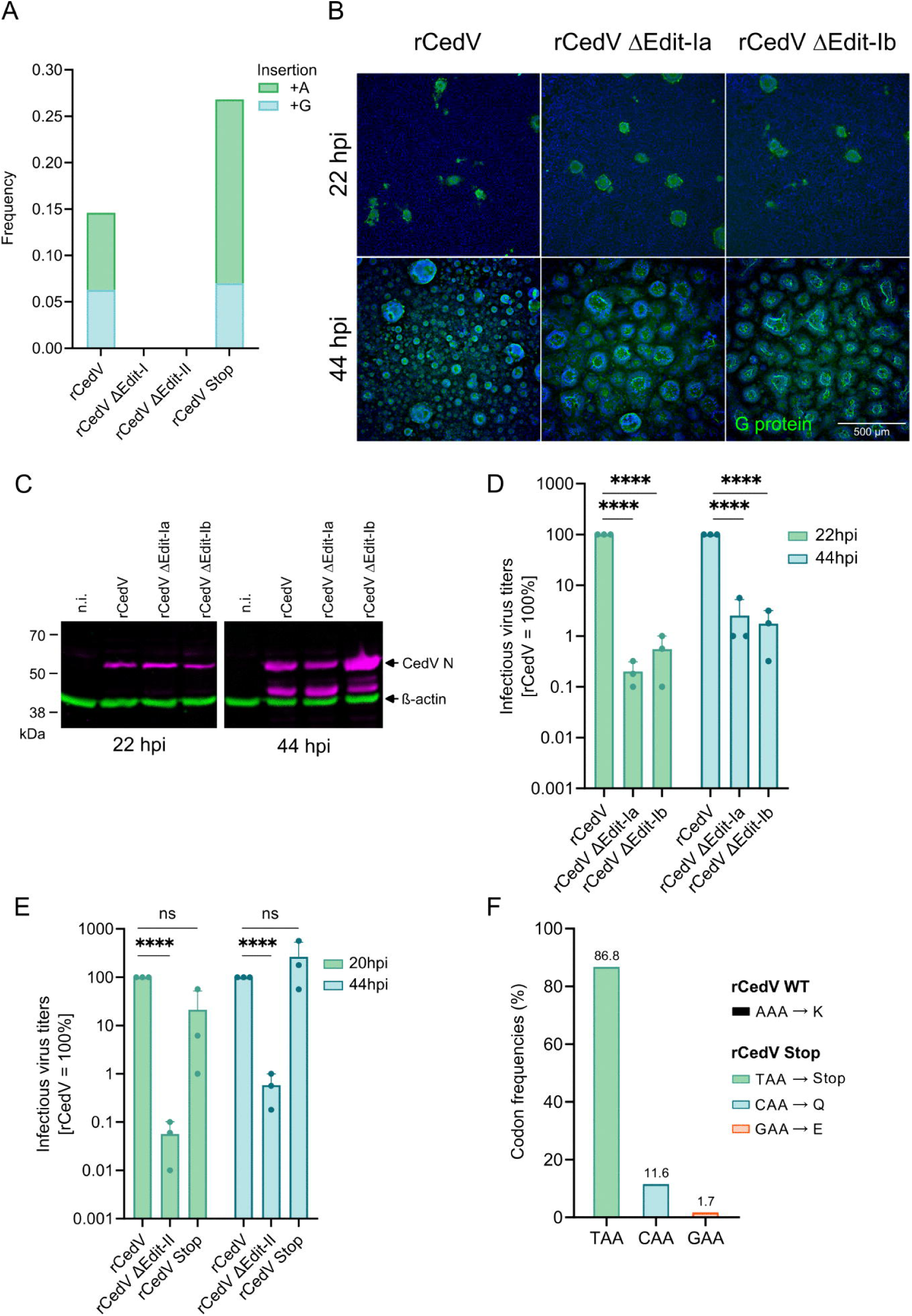
CedV P mRNA editing is required for efficient release of infectious virus. **(A)** Frequencies of adenosine and guanosine at editing nucleotide position 3312 for rCedV, rCedV ΔEdit-I, rCedV ΔEdit-II, and rCedV Stop. Frequencies of A and G insertions [1 =100%] are indicated by green and blue, respectively **(B)** Detection of rCedV ΔEdit-I and rCedV in BSR-T7/5 cells at 22 and 44 hpi by immunofluorescence with two independently rescued rCedV ΔEdit-I (Ia and Ib) viruses (green: G, blue: Hoechst 33342 stain). **(C)** Western blot detection of the CedV N protein (magenta) to confirm comparable infection and virus protein expression after infection at an MOI of 0.1. **(D)** Infectious virus titers in the supernatants of infected cells at 22 and 44 hpi. The titers of rCedV were set to 100% for each experimental replicate (n=3). **(E)** Infectious virus titers in the supernatants of rCedV, rCedV ΔEdit-II-and rCedV Stop (MOI 0.01)-infected cells at 20 and 44 hpi. The titers of rCedV were set to 100% for each experimental replicate (n=3) **(F)** Frequencies of the mutations in the inserted stop codon of the rCedV Stop at 20 h post transfection. The data (in d–f) are presented as the mean ± SEM. Statistical analysis was performed using one-way ANOVA, followed by Dunnett’s multiple comparisons test. Statistical significance is indicated as **** (p < 0.0001). Each dot indicates one independent experiment.

Despite comparable infections at 22 hpi, as assessed by immunofluorescence (Fig. 8B) and by western blot detection of CedV nucleoprotein N (Fig. 8C), the titers of infectious virus released into the supernatant were reduced by 505-fold (ΔEdit-Ia) and 181-fold (ΔEdit-Ib) relative to those of rCedV at 22 hpi (Figure 8D). Although the titers of both mutants increased by 44 hpi, they remained 39-fold (ΔEdit-Ia) and 57-fold (ΔEdit-Ib) lower than the wild-type levels, providing evidence that the loss of P gene editing, and consequently U protein expression, impaired the formation and/or release of infectious virions.

The negative impact of the editing site mutations in rCedV ΔEdit-I was confirmed by the rCedV ΔEdit-II mutant lacking synonymous downstream mutations. The infectious virus titers of rCedV ΔEdit-II decreased by 1793-and 177-fold at 20 and 44 hpi, respectively, at a lower MOI of 0.01 (Fig. 8E). Limitations in rCedV ΔEdit-II spread and larger syncytia sizes were shown by immunofluorescence imaging (S4 Figure 3). Notably, imaging analysis of rCedV Stop revealed smaller syncytia at 22 hpi, whereas more efficient spread was indicated by multiple infected cell syncytia at 44 hpi. This correlated with infectious virus titers in the cell culture supernatants, which reached wt rCedV levels at 44 hours post-infection (Fig. 8E). As the high infectious virus titers could be a result of either full functionality of the remaining 12 U amino acids in rCedV Stop or rapid acquisition of mutations that restore full U protein expression, the amplicon NGS sequences used for editing frequency calculation (Fig. 8A) were further screened for mutations in the U protein coding sequence. Indeed, the TAA stop codon present in rCedV Stop was mutated to CAA or GAA in 11.6% and 1.7% of the reads, respectively, leading to both the loss of the premature stop codon and the translation of a complete U protein (Fig. 8F). Given that wt rCedV comprises an AAA codon at that position, any contamination by authentic CedV virus or sequences could be excluded. The rapid selection of an escape mutation in rCedV Stop further demonstrates the major impact of U protein expression on CedV replication.

The effects of mRNA editing, and consequently U protein expression, on membrane dynamics at the cell surface were assessed by electron microscopy and immunofluorescence analysis of cells infected with rCedV or rCedV ΔEdit-I (Figure 9A). Compared with rCedV-infected cells, rCedV ΔEdit-I-infected cells exhibited larger vesicular structures containing G protein, as shown by immunofluorescence microscopy (Figure 9B). Furthermore, expression of Strep-tagged U in rCedV-infected cells revealed colocalization of U and the virus-expressed nucleoprotein within these vesicular structures (Figure 9C).

**Figure 9:**
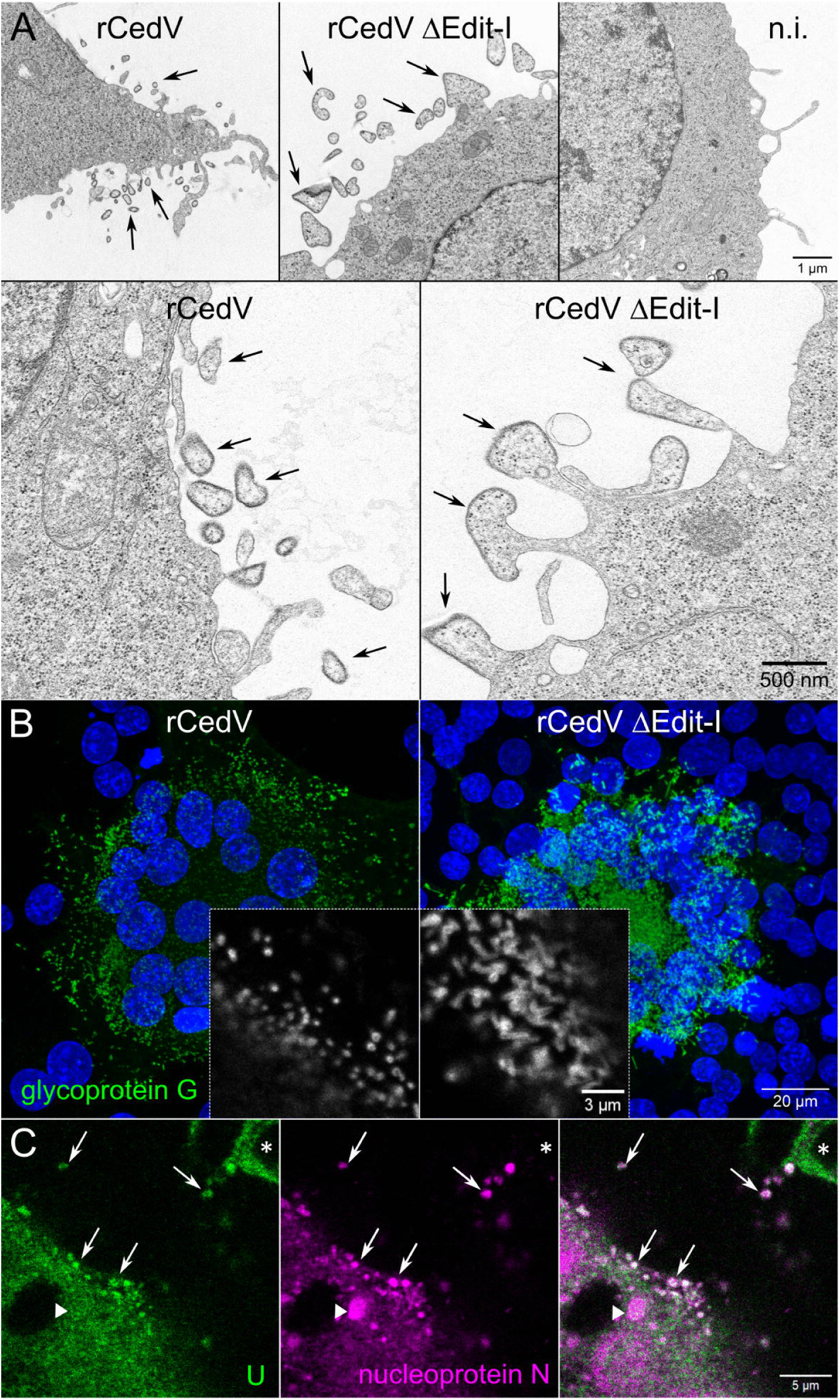
Enlarged plasma membrane-derived protrusions and vesicles in the absence of mRNA editing and localization of Strep-tagged U protein. (**A**) Ultrastructural analysis of plasma membrane protrusions and vesicles in BSR-T7/5 cells infected with rCedV or rCedV ΔEdit-I at an MOI of 0.05 and examined at 20 hpi. Arrows indicate infection-associated structures at the plasma membrane. n.i., non-infected. (B) Immunofluorescence detection of cell surface-localized glycoprotein G (green; displayed in grayscale in the magnified views). (C) Localization of plasmid-expressed Strep-tagged U protein in transfected BSR-T7/5 cells infected with rCedV and analyzed at 20 hpi. Arrows indicate vesicular structures containing colocalized U and N proteins; asterisks indicate non-infected cells expressing U; the arrowhead indicates a cytoplasmic CedV inclusion body.

### Attenuated replication of U protein-deficient CedV in IFNAR-KO mice

To assess the in vivo replication of the P gene editing–defective virus (rCedV ΔEdit-I), we used an interferon-α/β receptor knockout (IFNAR KO) mouse model, which is permissive for CedV infection [20]. Mice were infected intraperitoneally with 3 × 10^5^ TCID per animal, and viral loads were quantified by measuring CedV N gene RNA levels in selected tissues. At 3 days post-infection (dpi), N-gene RNA levels were reduced by 18-to 68-fold in rCedV ΔEdit-I-infected mice compared with those in rCedV-infected mice when all tested organs were considered. This demonstrated impaired replication of the editing-deficient virus in vivo. Differences between the viruses remained evident at 6 dpi, with 3-to 6-fold lower levels in the lung, kidney and spleen and 40-and 75-fold lower levels in the liver and lymph nodes than in the rCedV (Fig. 10). These results show that CedV P gene editing is necessary for efficient replication in vivo and that less efficient replication is detectable even in the absence of type I interferon signaling.

**Figure 10.**
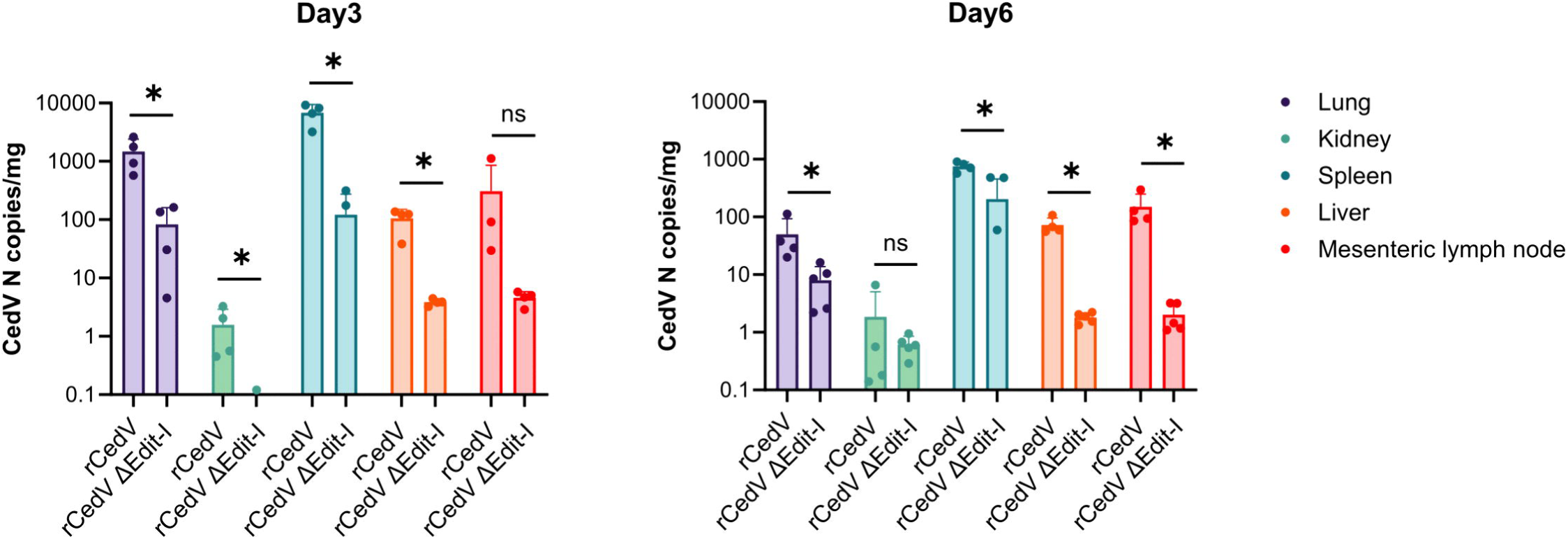
Attenuated replication of U-deficient CedV in IFNAR-KO mice. CedV N gene RNA levels in different organs after intraperitoneal infection at 3 x 10^5^ TCID per mouse. The data are presented as the mean ± SD. Statistical analyses were performed using GraphPad Prism. Differences between groups were analyzed using the Mann–Whitney U test. Statistical significance is indicated as * (p < 0.05); ns = not significant (p ≥ 0.05). Each group consisted of 4–5 animals.

## Discussion

Here, using CedV as an example, we provide evidence for noncanonical P-gene mRNA editing in a henipavirus and identify an alternative P-gene product, designated U, that displays selected sequence-, structure-, and localization-related similarities to henipavirus V and W proteins. Because this editing product is generated from an alternative P gene editing site and lacks the cysteine-rich region characteristic of canonical paramyxovirus V proteins, we refer to it as U protein. Strikingly, in contrast to most paramyxovirus V proteins, the CedV U protein is required for efficient virus rescue and infectious virus production. P gene editing-defective mutants yielded markedly reduced infectious titers, up to ∼1800-fold. A substantial effect of U on membrane curvature, with potential consequences for virus assembly, is further supported by the electron microscopy and immunofluorescence data. In the absence of U, enlarged plasma membrane-derived protrusions and vesicular structures were observed, suggesting that U contributes to the regulation of membrane remodeling during the viral life cycle. Together, these data point to a previously unrecognized contribution of a henipavirus P gene mRNA editing product to late stages of the CedV replication cycle.

Canonical paramyxovirus P gene editing is associated with the exclusive insertion of nontemplated guanosine residues at defined editing sequence motifs (e.g., AAAAGGG in NiV/HeV, mRNA orientation). In contrast, CedV editing occurs at a different upstream site (AAAAAAAG; mRNA orientation) and results in site-specific insertion of a single adenosine or guanosine residue. To our knowledge, this combination of editing site sequences and adenosine insertion has not yet been described for paramyxoviruses, expanding the concept of P gene editing beyond the previously established canonical motifs. Consequently, prevailing models of CedV gene expression, particularly the widely held notion that CedV lacks P gene mRNA editing, require revision. Importantly, we demonstrate the existence and functional relevance of this editing event in both infected cell cultures and a susceptible mouse model.

The apathogenic phenotype of CedV has been attributed primarily to altered receptor usage and a reduced capacity to antagonize innate immunity. CedV uses ephrin B1 and B2 for entry, whereas NiV and HeV preferentially use ephrin B2 and B3, which are thought to shape tissue tropism and host range [15]. CedV has been reported to lack canonical P gene editing and therefore to not express the classical V and W proteins implicated in host response antagonism [4]. Our findings challenge this concept by demonstrating the expression of an editing-derived U protein. However, this U protein lacks a cysteine-rich zinc-finger domain (ZnFD) broadly conserved in paramyxoviruses [21]. Loss of ZnFD could contribute to attenuation, whereas other conserved V and W protein-associated activities may be retained to support replication. In this context, U protein expression may be particularly important for time-limited spread in animal models, as previously observed [18], and for maintenance in its natural Pteropus bat reservoir. CedV mutants defective in P mRNA editing replicated less efficiently in vivo, supporting the need for the U protein in the infected host.

Previous work has shown that ablation of NiV V protein expression converts highly pathogenic NiV into a nonlethal virus in vivo [22]. However, whether the absence of a ZnFD in CedV is functionally equivalent to complete loss of the V protein in NiV, and whether this difference accounts for the interferon sensitivity and nonlethal phenotype of CedV, remains unsolved. It also remains to be determined whether the contribution of the CedV U protein to virus release, described in this study, reflects a broader, conserved function of V proteins in henipaviruses.

Mechanistically, the CedV editing site differs from conserved paramyxovirus motifs, which contain U-and C-rich nucleotide stretches on the template and promote exclusive G insertion via polymerase back-slipping [23]. In CedV, A or G insertions at the template sequence (U_7_C) imply a different mode of stuttering. Multiple A insertions are hallmarks of poly(A) tail formation by negative-strand RNA virus (NNSV) polymerases through slippage at U-rich transcription stop signals [24]. One possibility is that adenosine insertion at the CedV U-heptamer is driven by polymerase slippage at the CedV U_7_C signal, similar to Ebola virus glycoprotein mRNA editing, where adenosine insertion occurs at a U_7_G template sequence [25]. G insertions in the CedV P gene mRNA could be facilitated by the downstream template C (U_7_C) and the formation of a G:U wobble pair after a one-nucleotide backslip. A template AA dinucleotide upstream of the U-heptamer (AAU_7_C) could disfavor extended backsliding by creating A:A mismatches, which are not tolerated at canonical editing sites [23]. This model is also supported by structural constraints of NNSV polymerases, which form relatively short RNA:RNA hybrids with templates 7 to 9 bp in length in paramyxoviruses [26, 27]. The noncanonical U_7_C CedV editing site described here fits well in such a hybrid, which may be a structural prerequisite of polymerase slippage at this editing site.

Paramyxovirus P proteins are modular, comprising alternating disordered and ordered regions. Structural work on NiV indicates that the P protein multimerization domain forms a tetrameric coiled-coil and engages the polymerase in a distinctive tetrameric arrangement [28]. In CedV, we found the noncanonical editing site in a region analogous to the canonical NiV and HeV editing position, just upstream of the L-interacting domain. The differences in the intracellular localization in CedV-infected cells and the robust recruitment of P protein to cytoplasmic inclusion bodies versus the cytoplasmic and nuclear patterns of the U protein are consistent with different viral or host interaction partners.

AlphaFold predictions for CedV U and NiV V proteins indicate a conserved C-terminal domain architecture consisting of antiparallel β-sheets embedded within flexible regions and short helices. Such a β-sheet-containing CTD is broadly conserved among paramyxovirus V proteins and has been linked to the antagonism of MDA5-mediated signaling and interaction with DDB1 [21, 29].

Although we did not elucidate the effects of CedV U protein on individual innate immune pathways in detail, our data indicate that CedV U protein activity is antagonistic to type I interferon induction via RIG-I and TBK1-linked signaling. Whether the different C-termini of CedV U and P proteins contribute to antagonistic activities remains to be investigated. Global suppression of the interferon response was not pronounced under our experimental conditions. This may be related to the absence of the canonical cysteine-rich zinc-finger domain and suggests that the CedV U protein may possess functions beyond interferon antagonism.

In addition to having weak protein sequence similarities to those of the NiV and HeV V proteins, the CedV U protein presents features reminiscent of those of the NiV W protein. Whereas NiV P and V proteins are predominantly cytoplasmic, W protein is located in the nucleus and modulates host signaling through interactions with 14-3-3 proteins [11, 12]. In CedV, the U protein is localized to both the cytoplasm and the nucleus. Motif analysis predicts a bipartite nuclear localization signal generated within the edited region and candidate linear motifs compatible with 14-3-3 binding in the elongated C-terminus. By confirming the functional relevance of the editing-dependent NLS and demonstrating nuclear shuttling of U, our findings support a model in which CedV U displays selected V-and W-like properties. However, it remains unclear whether these similarities are either the result of a common evolutionary origin or of independent evolution. Experimental dissection will be required to test these interactions.

Genetic analysis further revealed the functional relevance of the U protein. Editing-defective viruses (rCedV ΔEdit-I and rCedV ΔEdit-II) and a mutant expressing a CTD-truncated U protein (rCedV Stop) demonstrate that U expression is critical for efficient infectious virus production. This requirement appears to differ from that of NiV, where the V protein can be ablated and the virus remains recoverable without trans-complementation [22]. Whether the U protein encodes a genuinely distinct function or whether related functions have been underappreciated in other henipaviruses remains to be determined. Nevertheless, our data suggest an unanticipated role of the CedV U protein in virion release, challenging the prevailing view that P protein variants act solely as immune modulators.

Only a few precedents exist for paramyxovirus P gene products that directly promote assembly or budding. With respect to NiV and Sendai virus, their C proteins facilitate virus release via ESCRT interactions, and the V protein of human parainfluenza virus type 2 counteracts tethering to promote virion release [30–32]. In this context, CedV U protein may likewise support efficient assembly or budding. Candidate tryptophan-dependent motifs predicted to interact with kinesin light chain 1 (KLC1) and the δ subunit of the COPI coatomer could contribute to transport or trafficking steps relevant to assembly and direct or indirect potential tethering inhibition.

Together, our findings reveal that the CedV U protein cannot be viewed as a canonical nonstructural interferon antagonist. Instead, the data support a model in which an editing-dependent U protein contributes to productive virion formation while immune-modulatory properties might have been lost. These results have two implications. First, these findings suggest that henipavirus P-gene products may influence processes associated with viral assembly and release more strongly than previously appreciated, motivating further investigation of mRNA editing related virus proteins across henipa-and related paramyxoviruses. Second, the discovery of noncanonical P gene editing in CedV highlights that genome annotation and risk assessment of emerging paramyxoviruses should explicitly consider alternative editing sites capable of generating protein variants in alternative frames, which may influence viral fitness in reservoir hosts and zoonotic potential.

## Materials and Methods

### Ethics Statement

The in vivo protocol was evaluated and approved by CECCAPP (Comité d’Evaluation Commun au Centre Léon Bérard à l’Animalerie de transit de l’ENS au PBES et au laboratoire P4) ethical comittee (Ministry of Higher Education and Research number CEEA - 015), with approval number 202410281755760.

### Cells

BSR-T7/5 cells were cultivated as described previously [33]. HeLa cells were obtained from the Collection of Cell Lines in Veterinary Medicine (CCLV; FLI Riems) and maintained in RPMI-1640 supplemented with 10% fetal calf serum (FCS). Human neuroblastoma SK-N-SH cells (ATCC HTB-11, [34]) were obtained from LGC (UK) and cultured in ZB28 medium (Ham’s F12/IMDM, 1:1) supplemented with 10% FCS. The Pteropus giganteus cell line PATGV was described previously [35].

### Viruses and full-length cDNA clones

Recombinant Cedar virus (rCedV) based on the CedV isolate CG1a genome sequence (GenBank: NC_025351.1) and a red fluorescent reporter–expressing rCedV have been described previously [36, 37]. The recombinant viruses rCedV ΔEdit-I, rCedV ΔEdit-II and rCedV-Stop (cDNA clone mutagenesis described below) were rescued in BSR-T7-dStrepU cells by cotransfection of the full-length CedV cDNA plasmid and support plasmids, together with pCAGGS-T7Pol encoding a codon-optimized bacteriophage T7 RNA polymerase [38]. Plasmids were transfected at the following amounts per 3.5 cm well: pCAGGS-CedV-N (1.25 µg), pCAGGS-CedV-P (0.8 µg), pCAGGS-CedV-L (0.4 µg), full-length CedV cDNA plasmid (3.0 µg) and pCAGGS-T7Pol (0.5 µg) using Lipofectamine 2000 (Thermo Fisher Scientific) according to the manufacturer’s instructions. Viral genomes of all recombinant viruses were confirmed by next-generation sequencing (NGS) of virion RNA extracted from culture supernatants.

Working virus stocks of rCedV were generated by infecting ∼80% confluent SK-N-SH cells in T75 flasks with 1 ml of BSR-T7/5 rescue supernatant or preexisting virus stocks. Stocks of rCedV ΔEdit-I, rCedV ΔEdit-II and rCedV-Stop were obtained from infected complementing BSR-T7-dStrepU cell cultures. Viral titers (TCID50) were determined by endpoint dilution titration on BSR-T7/5 and SK-N-SH cells.

### Construction of mutant full-length CedV cDNA clones

To generate pCedV-ΔEdit-I, 16.6-kb and 3.9-kb fragments were amplified from the CedV full-length cDNA clone using primer pairs 1/2 and 3/4, respectively (S7 Table 1), and assembled by hot fusion cloning [39]. The adenine heptamer at the noncanonical editing site was disrupted by A-to-G substitutions at CedV genome nucleotide (nt) positions 3307 and 3310. In addition, editing-irrelevant marker substitutions (A-to-G and G-to-A) were introduced at nt 3322 and 3342 to facilitate unambiguous identification of the mutant virus, including in the event of reversion at the editing site during amplification on noncomplementing cells.

pCedV-ΔEdit-II. pCedV-ΔEdit-II, containing A-to-G substitutions at nt 3307 and 3310, was generated from two PCR fragments (1.6 kb and 3.0 kb) amplified with primer pairs 5/6 and 7/8, respectively (S7 Table 1). The two fragments were joined by fusion PCR using the primers 6/8. The resulting 4.6-kb fragment was assembled together with a 16.6-kb XmaI fragment from the CedV cDNA clone to yield pCedV-ΔEdit-II by hot fusion cloning.

pCedV-Stop was generated using a comparable strategy with the 9/10 primer pair.

### Sequence verification and data availability

All plasmid constructs were verified by whole-plasmid sequencing (Eurofins Genomics, Germany). All the mutant virus stocks were sequence-confirmed by next-generation sequencing (NGS) (Eurofins Genomics, Germany). Viral genome sequences are available at Zenodo (DOI 10.5281/zenodo.18642052).

BSR-T7/5 cells stably expressing CedV Strep-U (BSR-T7-dStrepU) were generated by transfection with pCAGGS-dStrepU-Puro, encoding an N-terminally Strep-tagged CedV U protein and a puromycin resistance cassette. Following puromycin selection, single-cell clones were isolated by limiting dilution in 96-well plates and screened for Strep-U expression by fluorescence microscopy and western blotting using Strep-tag detection.

### MS analysis

Quadruplicate cultures of SK-N-SH cells were grown to 80–90% confluence and infected with rCedV at an MOI of 1. Mock-treated cells served as controls. Infection was carried out for 60 min at 37°C with 5% CO□. Following incubation, the inoculum was removed, and the cells were maintained in ZB28 medium (FLI biobank) until harvest. At 48 hpi, the cells were detached, transferred into 2 ml tubes and precipitated by centrifugation at 300 × g for 5 min (Fresco 21 [Thermo Scientific]). The cell pellets were resuspended in 200 µl of RIPA buffer (50 mM Tris/HCl (pH 7.5), 1% v/v IGEPAL CA-630, 0.5% v/v sodium deoxycholate, 150 mM NaCl in UP H2O) supplemented with protease inhibitors (1 mM pepstatin A [SERVA Electrophoresis GmbH], 1 mg/ml leupeptin [SERVA Electrophoresis GmbH], 100 mM PMSF [SERVA Electrophoresis GmbH]) and 0.5 M EDTA. The lysates were incubated on ice for 30 min, followed by vortexing and centrifugation at 10,000 × g (Fresco 21 [Thermo Scientific]) for 10 min at 4°C. The supernatants were transferred into new tubes, and glycerol (Carl Roth) was added to a final concentration of 10% (v/v). The protein concentration was determined by a BCA protein assay (Fisher Scientific) and measured on a GloMax Discover Microplate Reader (Promega). For proteomic analysis, 20 µg of total protein per sample was resuspended in 1× elution buffer (1× LDS [Thermo Fisher Scientific] supplemented with 100 mM DTT [SigmalAldrich]) and incubated at 70 °C for 10 min at 1400 rpm in a Thermomixer (Eppendorf). The samples were subsequently processed for mass spectrometry using a previously described in-gel digestion protocol [40]. A total of 400 ng of peptide per sample was separated on an Aurora Ultimate 25×75 C18 UHPLC column (ionopticks) using a nanoElute HPLC system (Bruker). Separation was conducted with a 100-min gradient at a constant flow rate of 300 nl*min^-1^ with increasing ACN concentration from 2% MS-grade ACN/0.1% formic acid (0–2 min) to 17% (2–67 min), 27% (67–88 min), 95% (88–97 min) and 4% (97–100 min). The column was mounted on a timsTOF HT mass spectrometer (Bruker) operated in standard PASEF data-dependent acquisition mode with a cycle time of 1.1 sec. Raw data were processed with MaxQuant (version 2.4.2.0) and searched against the SwissProt (42,338 entries) and Trembl (54,436 entries) (UP000005640, v20200117) *Homo sapiens* databases and all CedV proteins (JQ001776.1). To identify potential unknown viral open reading frames, the nucleotide sequences of all viral proteins were translated into all possible reading frames using the six-frame translation tool in MaxQuant (version 2.4.2.0), with a minimal AA length of 20. Downstream data analysis was performed using RStudio (version [2023.06.1]).

### Antibodies, sera and imaging

Polyclonal serum against CedV N was generated at the FLI by immunization of rabbits with a purified CedV N antigen from *Spodoptera frugiperda* (Sf9) insect cells infected with a recombinant baculovirus encoding the CedV N carrying an N-terminal histidine tag. CedV G–specific mouse monoclonal antibodies were generated at the FLI by immunization of mice with purified CedV G antigen produced in *Leishmania tarentolae*, as previously described by HeV G [41]. A Strep-tag–specific polyclonal rabbit antibody (Thermo Fisher Scientific, PA5-119772) diluted in PBS was used at 1:1,000 for immunofluorescence and 1:2,500 for western blotting. For immunofluorescence microscopy, Alexa Fluor 488– and Alexa Fluor 568–conjugated secondary antibodies (Thermo Fisher Scientific) were used at dilutions of 1:1,000 in PBS. For Western blotting, IRDye 680RD anti-mouse and IRDye 800CW anti-rabbit secondary antibodies were used (Li-COR Biosciences, Germany). Alternatively, rabbit anti-FLAG tag (Proteintech, Germany), mouse anti-β-Actin antibody (Proteintech, Germany) and their corresponding HRP-conjugated anti-Rabbit and anti-mouse secondary antibodies (Thermo Fisher Scientific) were employed for chemiluminescence detection. Nuclear DNA was counterstained with Hoechst 33342 (1 µg ml−1 in PBS). Fluorescence microscopy images were acquired with a Leica THUNDER imager DMi8 with 20×/0.40 dry HC PL FLUOTAR L objectives and LAS X software and with a Leica Stellaris 8 confocal laser scanning microscope with a 63 x/1.40 oil immersion HCX PL APO objective and LAS AF software (Leica Microsystems, Germany). Fluorescent Western blots were imaged with an Odyssey CLx infrared imaging system (Li-COR Biosciences). The images were processed with the Fiji (ImageJ 1.53t) software package [42].

Quantification of nuclear fluorescence intensities was performed using a custom automated image analysis workflow developed in Zeiss arivis Pro v.4.5.0. Confocal images were denoised using the Discrete Gaussian filter and segmented using the Blob Finder algorithm for nuclear identification and Huang thresholding for detection of U protein-positive regions. Nuclear and cytoplasmic compartments were defined based on the segmentation masks, and the mean fluorescence intensity of U protein signal was quantified separately in each compartment. Quantification data were exported and visualized using GraphPad Prism v.10.4.1. Statistical significance was assessed using ordinary one-way ANOVA.

### Electron microscopy

For transmission electron microscopy, cells were fixed in 2.5% glutaraldehyde in 0.1 M sodium cacodylate buffer, followed by post-fixation with 1% aqueous osmium tetroxide (OsO□). Samples were subsequently contrasted with 1% aqueous uranyl acetate and dehydrated through a graded ethanol series. After dehydration, the samples were transitioned into propylene oxide and embedded in an epoxy resin composed of glycid ether (Epon 812), dodecenylsuccinic anhydride (DDSA), methyl nadic anhydride (MNA), and DMP-30 as a polymerization accelerator. Resin polymerization was carried out at 60 °C for 3 days. Ultrathin sections were collected on glow-discharged nickel grids and sequentially contrasted with 2.5% uranyl acetate in 25% ethanol and aqueous lead citrate. Samples were examined using a Talos F200i transmission electron microscope operated at an accelerating voltage of 80 kV.

### RNA preparation, reverse transcription, PCR and amplicon sequencing

Total RNA was isolated from HeLa and SK-N-SH cells at 48 h post infection (hpi) using an RNeasy Mini Kit (Qiagen, 74104) according to the manufacturer’s instructions. First-strand cDNA synthesis was performed with a RevertAid First Strand cDNA Synthesis Kit (Thermo Fisher Scientific, K1622) using either oligo(dT) primers or the P gene–specific negative-sense primer 11 (S7 Table 1).

A 1,278-bp fragment spanning the CedV P gene was amplified from cDNA using primers 11 and 12 (S7 Table 1) and Phusion Hot Start Flex DNA Polymerase (New England Biolabs, M0535L). The PCR conditions were as follows: 94°C for 2lmin; 29 cycles of 94°C for 15ls, 58°C for 15ls and 72°C for 1lmin; 72°C for 10lmin; and a hold at 4°C. Amplicons were purified using the QIAquick Nucleotide Removal Kit (Qiagen, 28306) and subjected to Illumina-based amplicon sequencing (Eurofins Genomics). For PATGV cells, total RNA was extracted using the NucleoSpin RNA Mini kit (Macherey-Nagel). The P gene amplicon was generated as described above with P-specific primers and the SuperScript IV One-Step RT–PCR System (Thermo Fisher Scientific).

Sequencing data were processed in Geneious Prime using paired-end read assembly, trimming with BBDuk, read normalization (target coverage 40; minimum depth 6) and mapping to the CedV P coding sequence. Nucleotide insertions were called by SNP/variant analysis using a 3% frequency threshold and a minimum coverage of 10 reads.

### Cloning of P gene sequences into expression plasmids

A 2.2-kb fragment encompassing the CedV P gene coding region was amplified from the total RNA of rCedV-infected SK-N-SH cells by reverse transcription and PCR amplification using negative-sense primer 13, followed by PCR with primer 13/14 (S7 Table 1). The amplicon was inserted into a pCAGGS-based mammalian expression vector [43] carrying an N-terminal double Strep-tag (dStrep) by assembly with the 4.8-kb NotI/XhoI backbone fragment. Plasmids encoding the unedited P open reading frame were designated pCAGGS-dStrepCedV-P. Plasmids containing a single-nucleotide insertion at the P gene editing site were designated pCAGGS-dStrepCedV-U. To generate pCAGGS-dStrepU-Puro, a pCAGGS derivative containing an EMCV internal ribosome entry site (IRES) and a puromycin resistance cassette downstream of the EcoRI site was linearized with EcoRI. A 1.6-kb insert was amplified from pCAGGS-dStrepCedV-U using the primers 15/16 and assembled into the linearized backbone by hot fusion cloning.

### Plasmid DNA transfection

For the expression of cloned P gene cDNAs, virus-infected or non-infected cells were transfected using polyethylenimine (PEI), as described previously [44]. Transfections were performed in 3.5 cm dishes seeded with 5.0 × 10□ cells, using 6 µg of plasmid DNA per dish. **Leptomycin B treatment.** CRM1-dependent nuclear export was inhibited using leptomycin B (LMB; Sigma-Aldrich). At 20 h post-transfection, LMB was added to the culture medium of BSR-T7/5 cells at a final concentration of 40 ng mL⁻¹. After 3 h of treatment, the cells were fixed and processed for indirect immunofluorescence analysis.

### Protein structure analysis

Protein structure predictions were generated with ColabFold version 1.5.5 [45, 46] using its default parameters and the integrated Amber module for relaxation and energy minimization [47]. A total of five models per protein were predicted and ranked according to their mean pLDDT. Structural evaluation and visualization were performed with ChimeraX [48].

### IFNβ promoter and interferon-stimulated response element (ISRE) luciferase reporter assays

For human IFNβ promoter induction assays, HEK293T cells were transfected with an IFN-β promoter firefly luciferase reporter plasmid (Promega, USA) together with other plasmids as indicated in the corresponding experiments using PEI 25 kDa transfection reagent (Polyplus, France). At 24 h post transfection, the cell lysates were mixed with the substrate D-luciferin. For the ISRE luciferase assay, HEK293T cells were transfected with the ISRE reporter plasmid (#90402; Addgene, USA) as indicated. At 24 h post transfection, the cells were treated with universal IFN-I (800 U/ml) overnight, and the cell lysates were mixed with the substrate D-luciferin. Luminescence was measured with a TECAN Spark 3 microplate reader (Tecan Group Ltd., Switzerland).

### Infection of mice

All work involving live CedV infections was performed under BSL-2 conditions at PBES, Lyon, France. IFNAR-/-mice were inoculated intraperitoneally (IP) with 3.10^5^ TCID50 of rCedV or rCedV ΔEdit-I (volume inoculated of 200 µl) under inhalational anesthesia with isoflurane (isoflurane: ISOFLUTEK 1000 mg/g; Laboratorios Karizoo S.A.; anesthesia station: MiniHUB-V3; TEM SEGA, France). After CedV IP infection, the mice were monitored daily, and selected organs (lungs, kidneys, spleen, livers and mesenteric lymph nodes) were collected at the designated time points (3 and 6 days post-infection). The animal groups were as follows: rCedV-wt_d3: 4 animals; rCedV-wt_d6: 4 animals; rCedV-DelEdit_d3: 4 animals; and rCedV-DelEdit_d6: 5 animals.

### Sample collection and RT-qPCR

The mice were euthanized by cervical dislocation under isoflurane. The lungs, spleen, liver, kidneys and mesenteric lymph nodes were harvested. The tissues were snap-frozen at −80°C for molecular analyses. For RNA extraction, ≤30 mg of tissue was homogenized in RA1 buffer (NucleoSpin RNA, Macherey-Nagel) containing β-mercaptoethanol using a MagNA Lyser and centrifuged, and the supenatant was mixed with 70% ethanol before column-based purification according to the manufacturer’s instructions. RNA was used directly for viral detection or DNase treatment. RT-qPCR was performed using the Luna Universal One-Step RT-qPCR Kit on a StepOnePlus system. Viral loads are expressed as copies/mg of tissue. CedV-N was detected using the specific primers 17 and 18.

## Author contributions

Conceptualization: SF, FB, BH, OR. Investigation: HS, PS, IR, GP, LR. Formal Analysis: RK, DSU, HS, PS, FB. Resources: SD. Visualisation: RK, DSU, HS, PS, FB. Supervision: SF, FB, OR, BH. Writing – Original Draft Preparation: SF, HS, PS. Writing – Review & Editing: all authors.

## Acknowledgments

We are grateful to Angela Hillner, Institute of Molecular Virology and Cell Biology, for perfect technical assistance. We thank Sven Sander and Sven Reiche for support with the production of monoclonal antibodies, and Mandy Jörn and Kristin Vorpahl for Electron Microscopy support. This work was supported by intramural funding to Stefan Finke. We acknowledge the contributions of the CELPHEDIA Infrastructure (http://www.celphedia.eu/), especially the center AniRA in Lyon. We acknowledge the contribution of SFR Biosciences (Universite Claude Bernard Lyon 1, CNRS UAR3444, Inserm US8, ENS de Lyon) and notably the PBES facility. Ilona Ronco was funded for her Ph. D by the Direction Générale de l’Armement (DGA) and the Agence Innovation Défense (AID).

## Data and Materials availability

The authors declare that the data are available in the publication, supplementary materials or at the public depository Zenodo (DOI 10.5281/zenodo.18642052).

## List of Supporting Information

**S1_Rawgel. Original uncropped and unadjusted Western Blot images.**

**S2 Figure 1.**
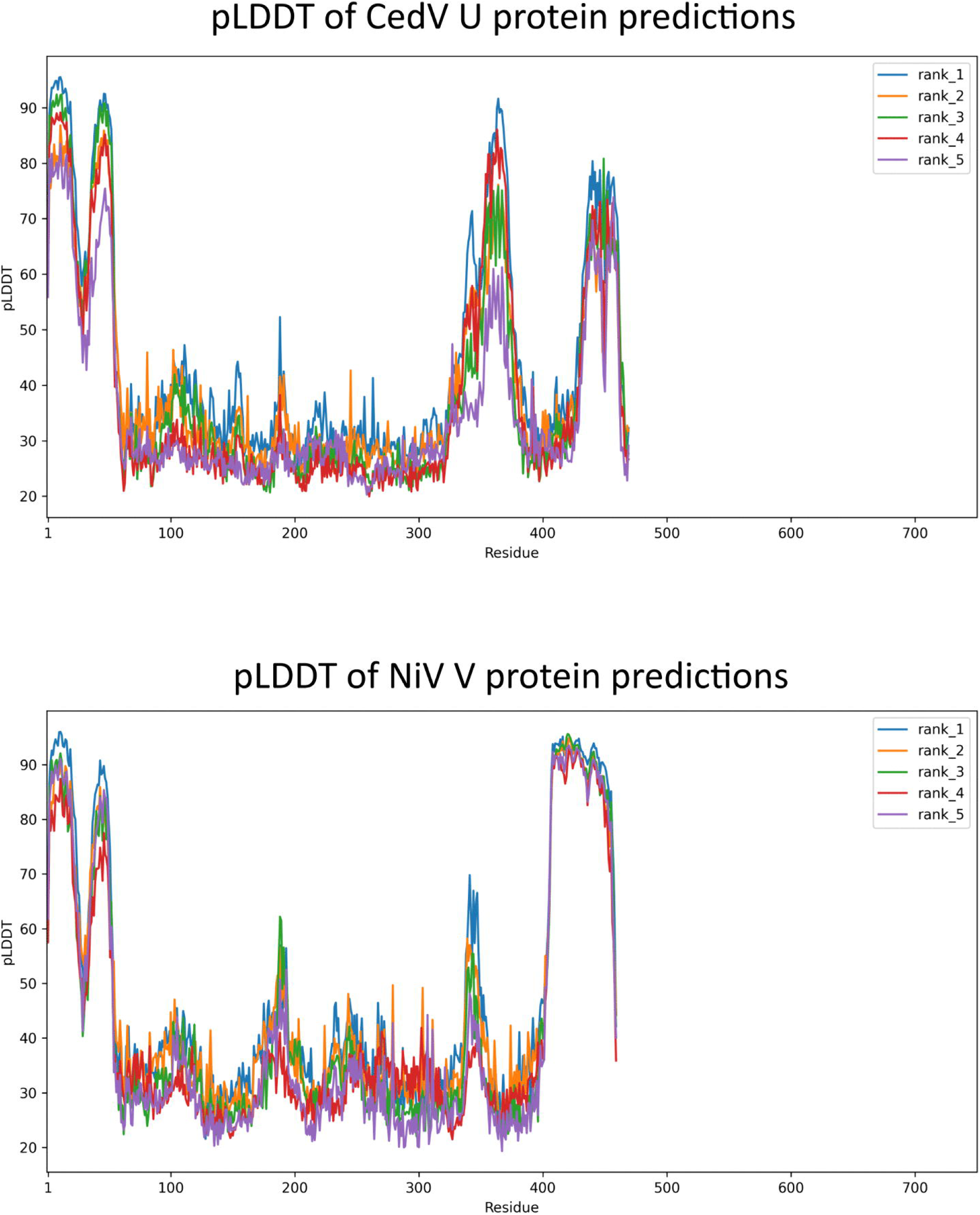
Local distance difference test plots (pLDDTs) indicating the prediction accuracies of the IDRs and structured domains in CedV P and NiV. **V.** High accuracies were determined for the U CTD (residues 402-469) and V CTD (residues 408-459).

**S3 Figure 2.**
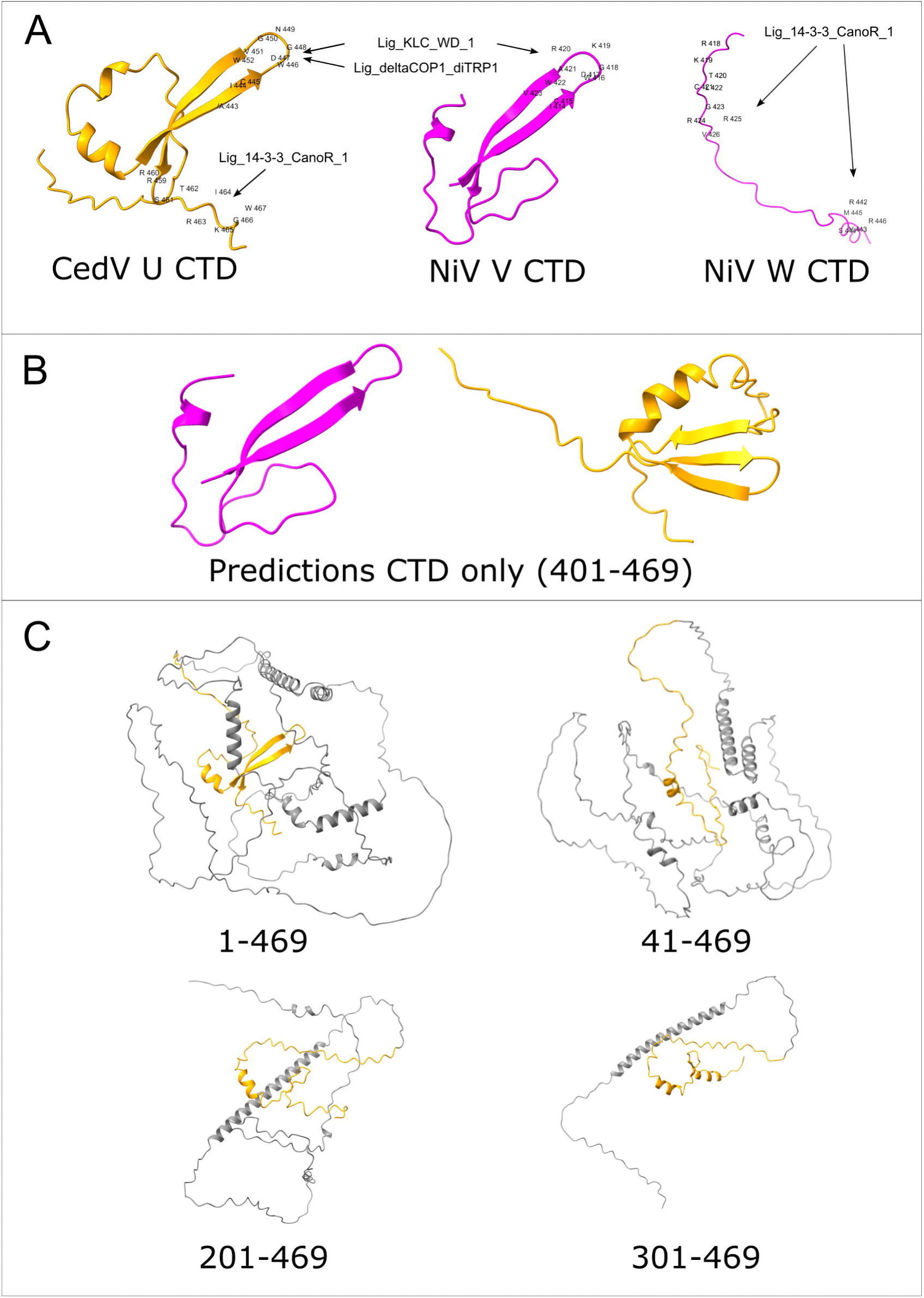
ELM linear motif predictions and structure predictions for partial U protein sequences. (A) ELM linear motif search revealed a potential Lig_KLC_WD1 motif in both the beta-sheet loops of CedV and NiV while CedV also exhibited a Lig_deltaCOP1_diTRP1 motif in that region. A potential Lig_14-3-3-CanoR1 motif was predicted for the U proteins disordered C-terminus, while two Lig_14-3-3-CanoR1 were predicted for the disordered NiV W protein C-terminus (B) Structural predictions for the CTDs (in CedV U positions 401 - 469) only revealed robust prediction of the conserved paramyxovirus V protein structure in NiV V, while the CedV CTD exhibited a change to a three shorter beta-sheet structure. (C) Predictions for sequential N-terminal truncations suggest an important role of the N-terminal 41 amino acid for folding to the typical V protein ‘s C-terminal fold.

**S4 Figure 3.**
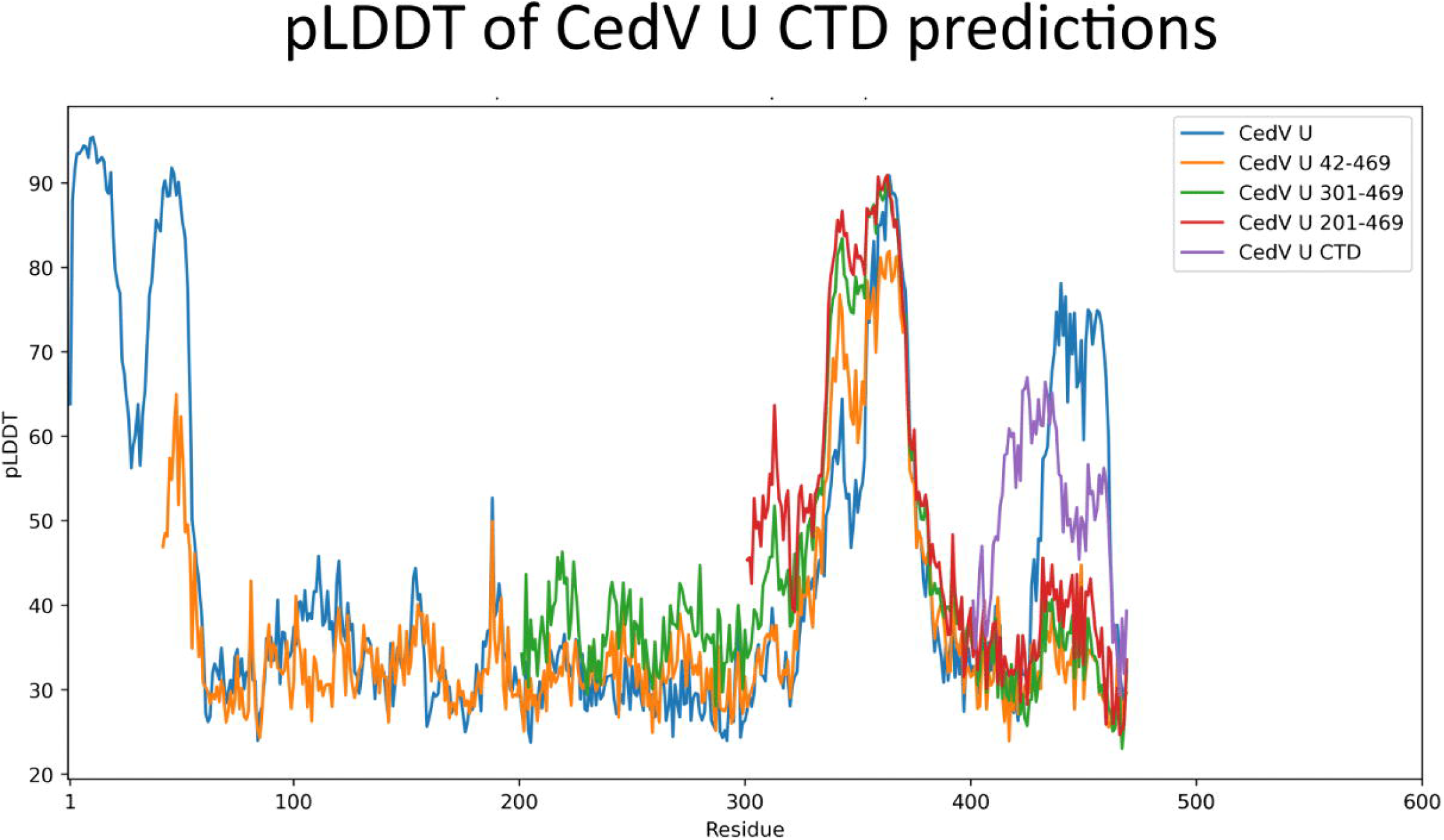
Local distance difference test plots (pLDDTs) indicating the prediction accuracies of the IDRs and structured domains in CedV U and NiV. **V.** Highest accuracies were determined for the U CTD (residues 402-469) in the complete U protein predictions (1-459; blue), followed by prediction for the CTD (402-469, purple). Analysis of N-terminally truncated U protein sequences beginning at residues 42, 201, and 301 revealed low pLDDT values.

**S5 Figure 4:**
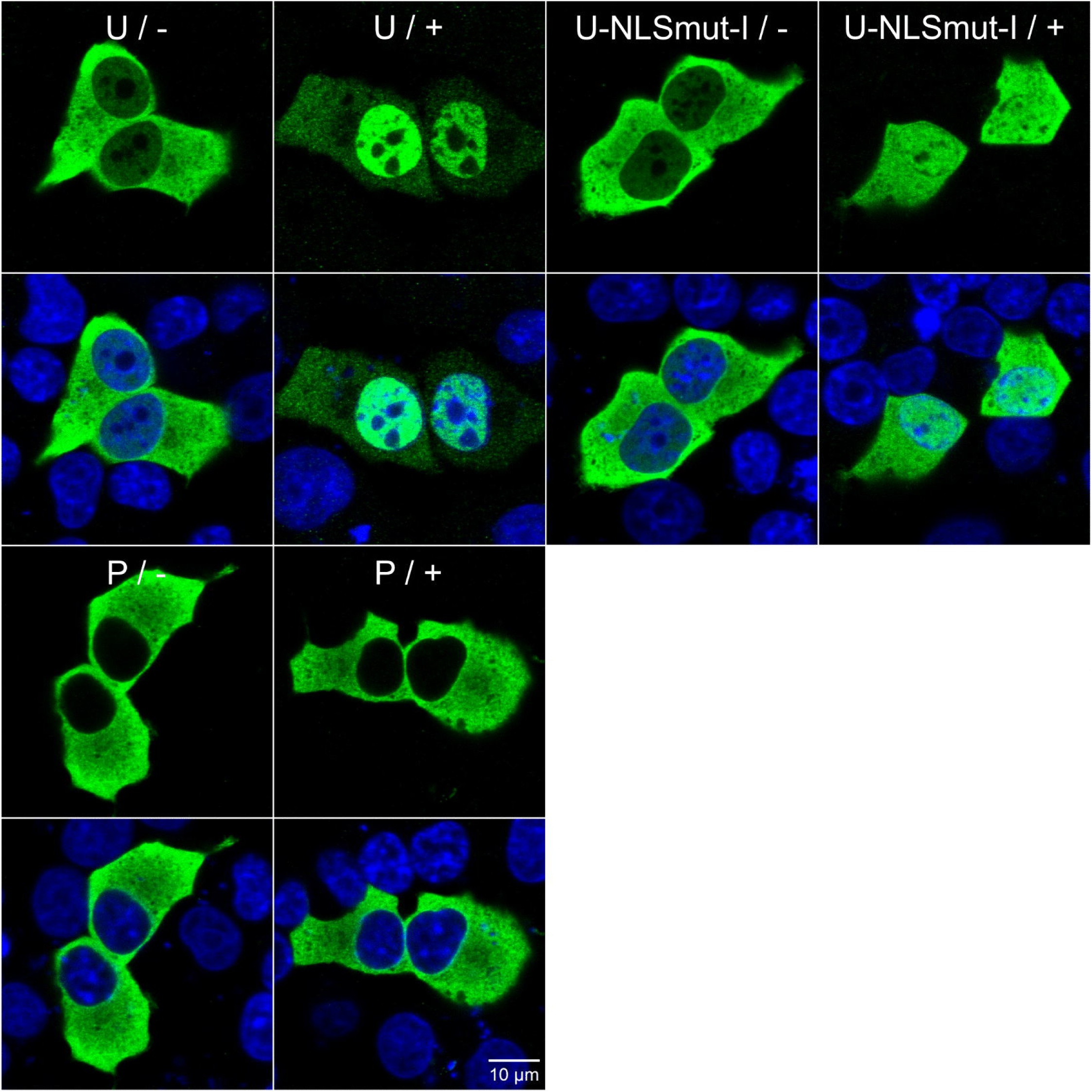
Nuclear shuttling of U and dependency on the predicted editing dependent NLS. Cells were transfected with expression plasmids for the indicated Strep-tagged proteins. 18 hrs later, leptomycin was added for 3 hrs. After immunofluorescence detection of the proteins (anti Streptag-antibody) and Hoechst 33342 stain, the images were acquired by confocal laser scan microscopy. Samples with (+) and without (-) leptomycin are compared.

**S6 Figure 5.**
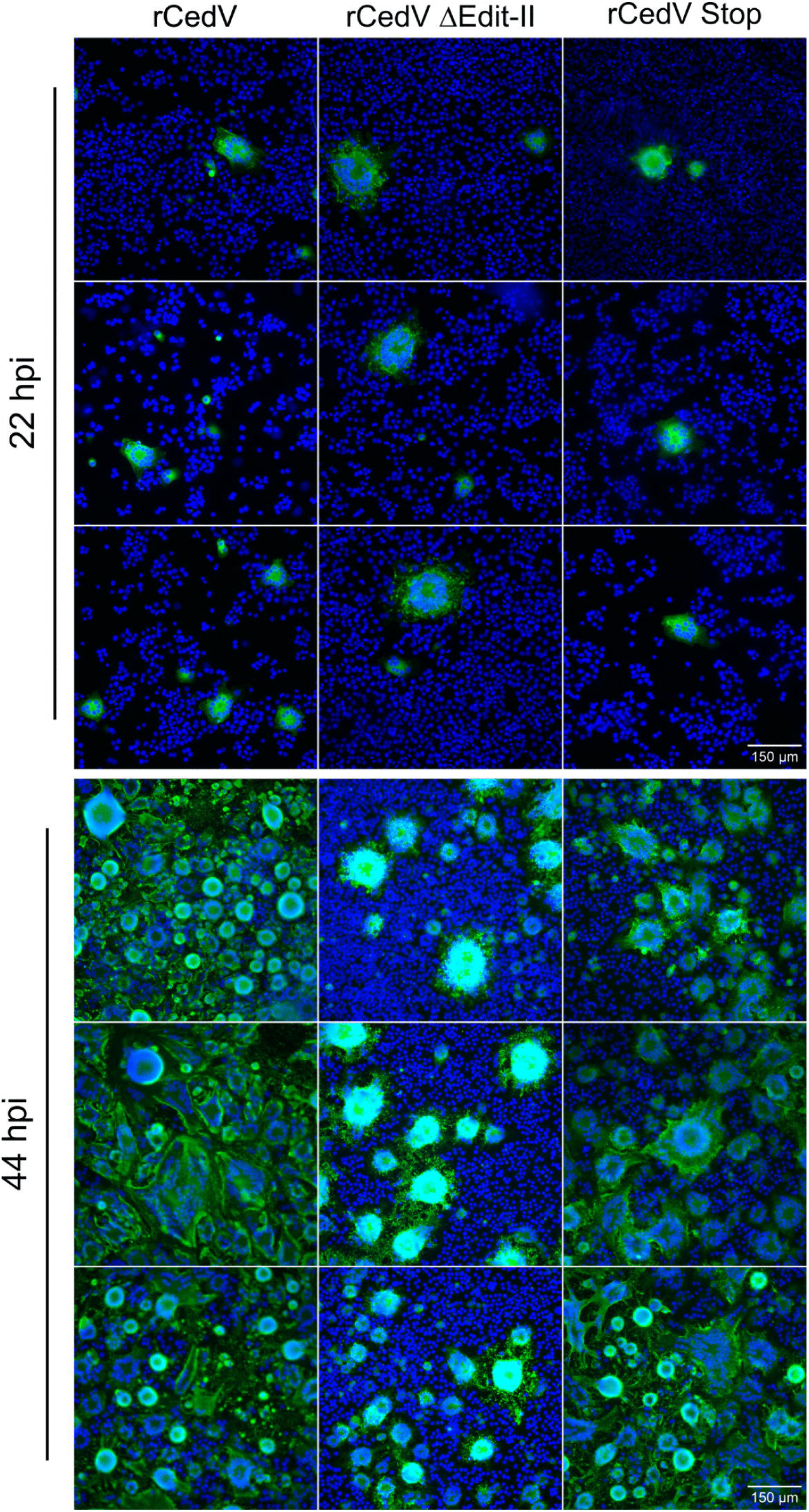
Infection of cells with rCedV ΔEdit-II and rCedV Stop at 22 and 44 hpi (MOI 0.01). Detection of rCedV ΔEdit-II, rCedV Stop and rCedV in BSR-T7/5 cells at 22 and 44 hpi by immunofluorescence (green: G; blue: Hoechst 33324). For each virus and timepoint, representative images are shown.

---

**S7 Table 1.**
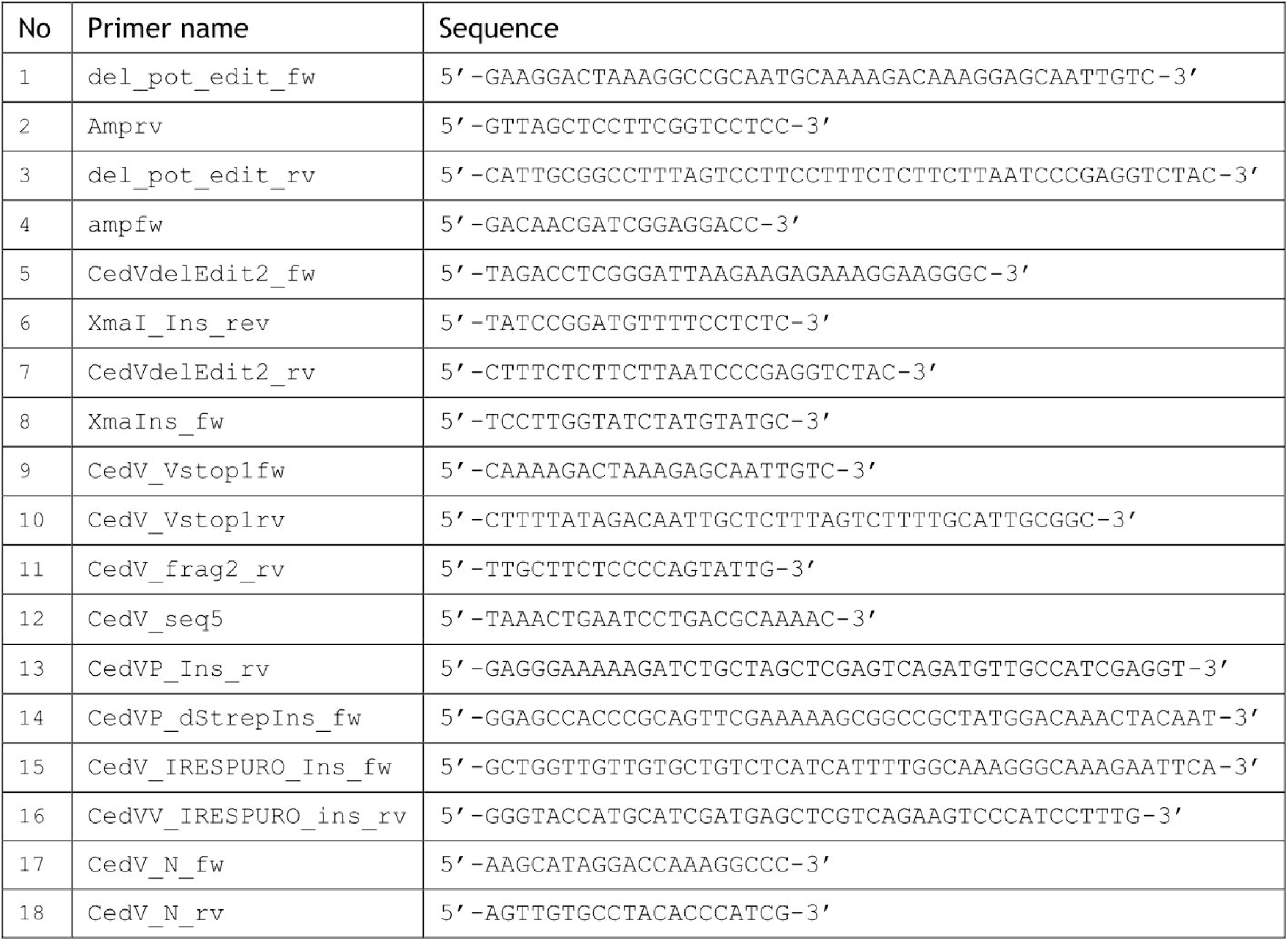
List of DNA-Oligonucleotide primers.

**S8 Table 2.**
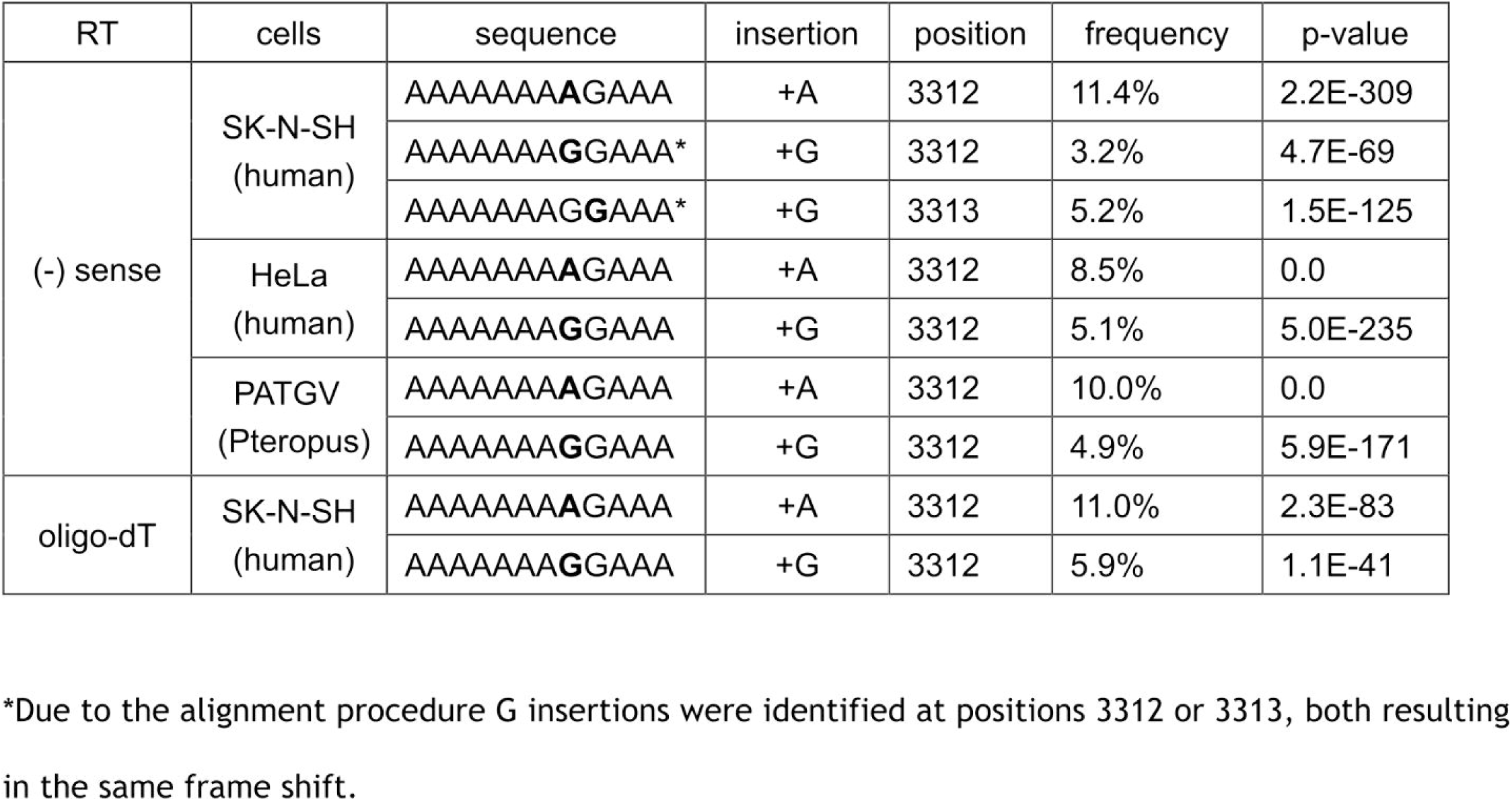
Amplicon sequencing revealed a hotspot of single-nucleotide insertion at CedV at position 3312/3313. (-) sense and oligo-dT reverse transcription (RT) primers were used to amplify P gene amplicons from RNA isolated from CedV-infected SK-N-SH, HeLa, and PATGV cells. The sequences around positions 3312/3313 with A or G insertions (bold letters) are indicated, and the frequencies of the respective insertions and the sum of all the 1 nt insertions per amplicon are indicated.

**S9 Table 3.** Amplicon sequencing of the rCedV, rCedV ΔEdit-I, rCedV ΔEdit-II, and rCedV Stop P gene sequences. NGS sequencing of P gene amplicons generated by RT-PCR ((-) sense RT-primers) from BSR-T7/5 cells at 24 hpi (MOI = 0.1). The frequencies of wt and single-nucleotide insertions at the noncanonical editing site at position 3312 are indicated.

| virus | Wt | +A | +G | editing frequency |
| --- | --- | --- | --- | --- |
| rCedV | 85.4 % | 8.3 % | 6.3% | 14.6 % |
| rCedV ΔEdit -I | 100 % | 0 % | 0 % | 0 % |
| rCedV ΔEdit -II | 100 % | 0 % | 0 % | 0 % |
| rCedV Stop | 73.2 % | 19,8 % | 7 % | 26.8 % |

